# Competition between bottom-up visual input and internal inhibition generates error neurons in a model of the mouse primary visual cortex

**DOI:** 10.1101/2023.01.27.525984

**Authors:** J. Galván Fraile, Franz Scherr, José J. Ramasco, Anton Arkhipov, Wolfgang Maass, Claudio R. Mirasso

## Abstract

The predictive coding theory, although attractive, is far from being proven. Supporters of this theory agree that bottom-up sensory inputs and top-down predictions of these inputs must be compared in certain types of neurons called error neurons. Excitatory neurons in layer 2/3 (E2/3) of the primary visual cortex (V1) are ideal candidates to act as error neurons, although how these error neurons are generated is poorly understood. In this study, we aimed to gain insight into how the genetically encoded structure of canonical microcircuits in the neocortex implements the emergence of error neurons. To this end, we used a biologically realistic computational model of V1, developed by the Allen Institute, to study the effect that sudden changes in bottom-up input had on the dynamics of E2/3 neurons. We found that the responses of these neurons can be divided into two main classes: one that depolarized (reporting positive errors; dVf neurons) and one that hyperpolarized (reporting negative errors; hVf neurons). Detailed analysis of both network and effective connectivity allowed us to uncover the mechanism that led to the dynamic segregation of these neurons. This mechanism was found to be the competition between the external visual input, originating in the thalamus, and the recurrent inhibition, originating mainly in layers 2/3 and 4. In contrast, we found no evidence of similar division and responses in excitatory infragranular neurons of layers 5 and 6. Our results are in agreement with recent experimental findings and shed light on the mechanisms responsible for the emergence of error neurons.

## Introduction

Since we began to interact with the environment, we learned to distinguish whether a movement is the result of our own action or due to the movement of objects external to us. To distinguish between these experiences, it is necessary to consider the effect that our actions have on incoming sensory information. The main framework accounting for this sensorimotor integration is **predictive coding**, which suggests that an internal representation of the world resides in the circuitry of the neocortex [1, 2]. This representation, which is used to make predictions about incoming sensory information, is continuously updated using perceived information from our environment. In particular, predictive coding theory suggests that E2/3 neurons in V1 compute the difference between bottom-up visual inputs (BU) and top-down predictions (TD) generated by higher cortical areas. As a result, the prediction errors computed by these neurons are transmitted to higher cortical areas to update the internal representation of the world. These prediction errors, which may come in two flavors, allow us to identify two classes of prediction-error (PE) neurons [1]: negative/positive prediction-error neurons (nPE/pPE) neurons that increase/decrease their firing rate relative to a baseline when sensory input is less/greater than predicted, and decrease/increase it in the opposite case. The appearance of prediction error neurons is thought to depend on the highly complex (and poorly understood) dynamics of the recurrent connections, which some theoretical studies suggest to encode features of the world in their weights distribution as a fast way to match sensory evidence [3].

Experimental evidence of prediction errors was found in many neural circuits, such as the visual cortex [4, 5], the auditory cortex [6], and the reward system [7]. While dopaminergic neurons have been found to signal mismatches between TD and BU inputs bidirectionally [8], E2/3 neurons in V1 can only signal mismatches unidirectionally, as the negative deviation of firing rates would be bounded from below due to the very low spontaneous firing rates exhibited by these neurons [9, 10]. Despite this, recent experimental studies using intracellular voltage recordings detected these negative deviations [5]. More recently, it was suggested that BU and TD inputs are integrated individually, without any sensorimotor expectation [11]. Consequently, the emergence of error neurons, as previously reported, might change radically and, therefore, could also be detected in the absence of TD signals, only as a response to BU signal features. However, stronger evidence for the former theory has recently been proposed, as the integration of individual inputs cannot explain the strong responses found when comparing stationary mice with moving mice during playback scenarios [12]. What is clear is that there is a segregation of the E2/3 neurons into two main and clearly distinguishable classes, one in which neurons depolarized (which we call dVf neurons) and another in which they hyperpolarized (which we call hVf neurons). This segregation appears to be essential for the emergence of PE neurons.

Therefore, understanding how the genetically encoded structure of canonical microcircuits (CMs) in the neocortex implements the emergence of error neurons remains an important open research question. The answer to this tangled question can be better elucidated from the implementation and analysis of computational models of CMs. Recent research has shed light into the computational roles played by different neuronal populations [13] and the conditions required for the emergence of PE neurons in V1 CMs [14]. In particular, a balance between excitation and inhibition in E2/3 neurons was suggested for these cells to act as PE neurons [14, 15]. However, when considering the V1 area, the role of neurons feature selectivity must be considered as a prominent feature in this computational mechanism, since some neurons have been found to act as feature detectors in many sensory pathways [16, 17].

Our aim in this study was to understand whether the mere presence of a time-varying BU visual input was sufficient to generate positive and negative error neurons in the V1 cortical layer of mice as suggested by previous experimental evidence [11], and if so, what caused the emergence of these two classes. We specially focused on the following aspects: the characteristics of the V1 connection network, the influence of neural and synaptic dynamics, and the competition between external and internal (recurrent) inputs. Although early studies on brain networks focused mainly on the structure and characteristics of the network [18, 19], numerous studies have shown that it is essential to include neural dynamics to have a correct description of the circuit behavior [20–22]. Actually, the fact that there exists an anatomical link between neurons does not guarantee that there is a functional connection, since this depends on the dynamic activity of the neurons. Likewise, the fact that functional connectivity exists does not imply that there is anatomical connectivity. In fact, functional hubs, that are not necessarily structural hubs, have also been identified as efficient nodes for information transmission in brain networks [23].

To account for the feature selectivity of V1 neurons, we used a biologically realistic model for the mouse area V1 and the lateral geniculate nucleus (LGN) of the thalamus that has recently been developed based on a large body of experimental evidences [24]. The use of this computational model, which we denote as the Billeh model (Fig. 1*A*), has several advantages: i) it considers neurons direction and orientation selectivity, which allows to explore how different configurations of BU input features affect the response of E2/3 neurons and compare them with experimental observations; ii) it gives direct access to the currents received by each neuron, which allows exploring the particular role of recurrent and BU inputs in the dynamics of the system; iii) it can provide insights on the role that inhibitory neurons play in the dynamics of the network.

**Fig. 1.**
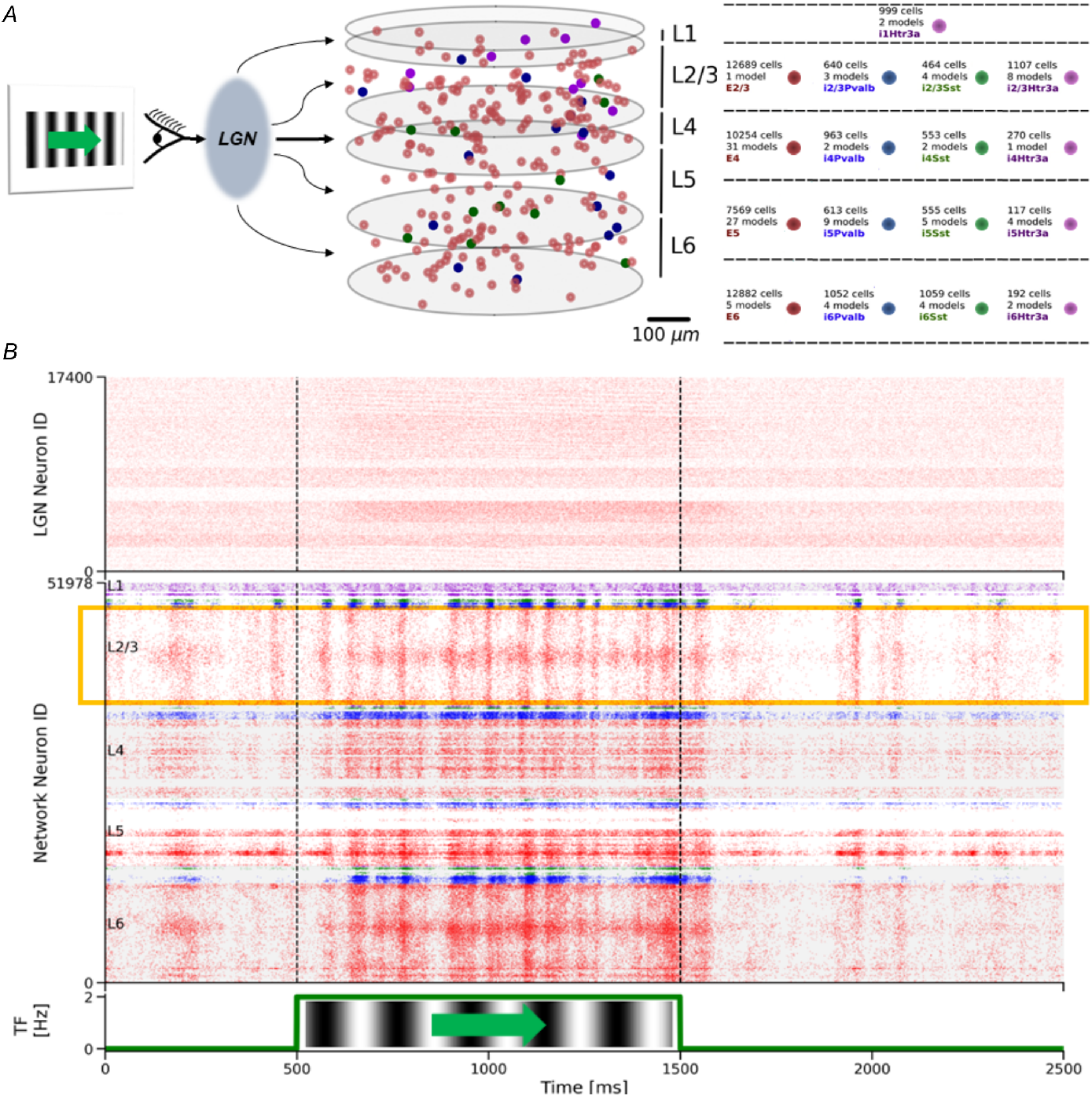
Billeh model and its response to the sudden onset of visual flow. *A*, The Billeh model describes a patch of mouse cortical area V1 and consists of 230,924 point-neurons divided into one excitatory and three main subclasses of inhibitory neurons (Pvalb, Sst, Htr3a). Each of these main classes are subdivided into various models of neurons. The V1 neurons in each population are evenly distributed within a cylindrical domain and within 5 laminar sheets: L1, L2/3, L4, L5 and L6. The cortical microcircuit model receives the BU input from a thalamus model that transforms the visual input into input currents. In this study we mainly focused on the detailed “core” region (400*μm* radius) of the model. This visualization shows only 0.5% of those core neurons. *B*, Top: Raster plot of the spike response of LGN units to visual stimulus. Middle: Laminar raster plot of the spike response of V1 neurons to the visual stimulus. Different colors represent different populations of neurons, following the same palette as in A. Several horizontal patches of higher firing rate can be seen due to the directional selectivity of individual neurons. The yellow box highlights E2/3 neurons, which are the main concern of this research. Vertical dashed lines indicate the period of visual flow. Bottom: Temporal frequency of the visual flow, which consisted of vertical gratings moving horizontally.

To gain insight into the emergence of dVf and hVf neuron classes, we performed extensive numerical simulations using the Billeh model to analyze the effect that different visual stimuli have on E2/3 neurons. To this end, we calculated the contributions of the different currents that reached each E2/3 neuron, whether excitatory or inhibitory, and determined the main sources of these currents. The aim was to explore the role that each type of neuron played in the cortical microcircuit. Our study revealed that E2/3 neurons exhibited a two-way response to visual flow perturbations, leading to two main classes of dVf and hVf neurons, similar to the behaviour observed for PE neurons under the influence of BU and TD inputs. This peculiar behavior was not observed in the infragranular layers (L5/6) which showed a one-way response to visual flow perturbations. The network dynamics analysis that we carried out allowed us to determine properties of the network that were not directly observed in its particular structure and to deepen our understanding of the computational properties of E2/3 neurons. In particular, we identified the layer 2/3 circuitry that enabled the presence of error neurons in the Billeh model, where also Parvalbumin neurons played a key role.

## Results

### Response of E2/3 neurons to a sudden onset of visual flow

Recent experimental studies, where visual flow and locomotion speed were coupled (closed-loop experiment) and sudden halts of visual flow (mismatches) were introduced, have identified depolarizing and hyperpolarizing excitatory neurons within L2/3 as nPE and pPE neurons, respectively [5]. Moreover, this segregation was also found when visual flow and locomotion were uncoupled (open-loop experiment), i.e. in the absence of explicit sensorimotor expectation, even if the mice remained stationary during the visual flow perturbation [5, 11]. Therefore, the absence of a locomotion-related TD input in the model should not qualitatively affect the behaviour of E2/3 neurons. Although locomotion is an important source of TD input to V1, it is certainly not the only one and, since the effect of the different TD inputs is not well understood, we focused our study on the effect that visual flow alone had on the organization of E2/3 neurons.

In our study we initially considered a visual BU input consisting of vertical static gratings that suddenly started to drift horizontally at a frequency of 2 Hz (see Methods for details and Fig. *1B*). In agreement with the bidirectional mismatch responses detected in previous experimental studies [5, 11], we found neurons with depolarizing and hyperpolarizing responses at the onset of visual flow (Fig. 2*A*). We classified each E2/3 neuron as **depolarized** (dVf) or **hyperpolarized** (hVf) with visual flow if its average input current change when visual flow is turned ON was greater than 0.05× rheobase current (dVf), or less than minus 0.05× rheobase current (hVf), being the rheobase current the threshold above which a neuron fires (see Methods for details and Fig. S1). E2/3 neurons that did not show a clear response to the stimulus were identified as **unclassified** (unc) (see Fig. S2). Before the onset of visual flow, the different components of the input current exhibited similar values for all classes of E2/3 neurons with an excitatory-inhibitory balance among recurrent currents, which resulted in an almost zero recurrent contribution, much smaller than the BU current (Table S1). However, once the visual flow was turned on, significant changes were observed between the different E2/3 classes. Notably, we found that depolarization of dVf neurons was primarily driven by an increase in BU currents, while recurrent inputs barely changed (Fig.*2A* and Table S1), maintaining the multi-pathway balance in the network, which was suggested as a requirement for the emergence of error neurons [14, 15]. Instead, the hyperpolarization of hVf neurons was caused by a reduction in recurrent excitation, which could have been caused by an increase in the activity of inhibitory presynaptic neurons, a reduction in the activity of excitatory presynaptic neurons, or a combination of both of them. In addition, we found a slightly larger number of neurons belonging to the hVf class as compared to the dVf class (dVf, 23.2% (2948/12689); hVf, 26.5% (3363/12689), unclassified, 50.3% (6378/12689)) (Table S1).

**Fig. 2.**
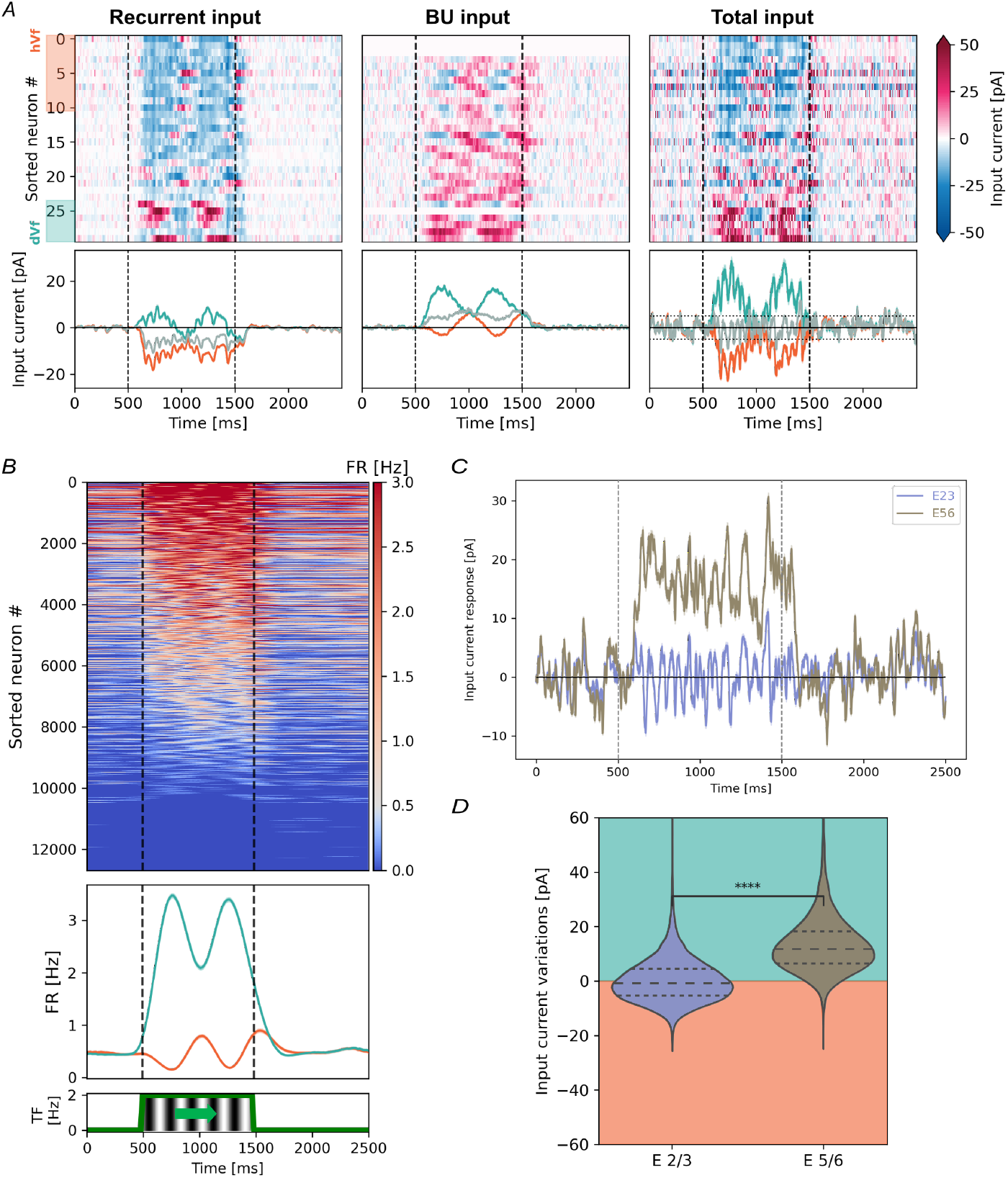
Diversity of E2/3 neural responses to sudden onset of visual flow. *A*, Average current responses of the different input current sources for each of the three classes of E2/3 neurons to the onset of visual flow with a spatial frequency of 2 Hz. Top: Heatmap for 30 randomly selected E2/3 neurons ordered by their average response. White rows indicate zero input current. Bottom: Average class traces. Horizontal dotted lines represent the classification threshold. Shading indicates the SEM measured across neurons and realizations. *B*, Top: Firing rates (FR) for the E2/3 neurons. Middle: FR averaged for dVf (turquoise) and hVf (orange) neurons. Bottom: Visual stimulus temporal frequency. *C*, Average populations trace of the input current response to the onset of visual flow. Shading indicates the standard error of the mean (SEM) measured across neurons and realizations. *D*, Probability distributions of the average input current responses of E2/3 and E5/6 neurons to the onset of visual flow (*t* = −110.7, *p* = 0, *n* = 33140, *n*_E2/3_ = 12689, *n*_E5/6_ = 20451, Welch’s t test). Turquoise and orange shades indicate regions of depolarization and hyperpolarization, respectively. The horizontal lines show the quartiles of the distributions.

Interestingly, the input current (Fig. 2*A*) and firing rate (Fig. *2B*) of dVf and hVf neurons had opposite phases, as if hVf neurons were silenced when dVf neurons became more active, and vice versa. We also found that certain baseline neural behaviours were significantly different between E2/3 classes. In particular, the mean and standard deviation of the baseline input current followed slightly different distributions for each E2/3 class (Fig. S3*A-B*). This suggested that the characteristic response of a neuron to the onset of visual flow is related to its baseline behaviour, which was also reported in experimental data [5].

### E5/6 and E2/3 neurons integrate visual input differently

Previous studies suggested that the infragranular layers, namely L5/6, integrate visual input in a different way as compared to L2/3 [1, 13]. In particular, the sudden onset of drifting gratings during an open-loop experiment triggered widespread depolarization within E5/6 neurons [5].

Our results obtained with the Billeh model receiving only visual input also showed a generalized depolarization of E5/6 neurons as a result of the onset of visual flow (Fig. 2*C-D* and Table S1), consistent with a positive integration of visual flow within this layer. In particular, E5/6 neurons exhibited a clear imbalance between depolarizing and hyperpolarizing responses (dVf, 62.7% (12816/20451); hVf, 0.3% (67/20451); unclassified, 37% (7568/20451)). In general, there was a significant difference between the average responses to the onset of visual flow in E2/3 neurons and E5/6 neurons (mean ± SD, L2/3: 0.2 ± 7.9 pA; L5/6: 13 ± 11 pA) (Fig. 2*D*).

### Connectivity differences in E2/3 classes

So far, the model has shown that E2/3 neurons can split into two classes in the sole presence of a time-varying visual input. In the following, we focus on unraveling the principles that lead to this effect. Previous computational studies on cortical microcircuits have shown that error neurons may require a balance between excitation and inhibition in multiple pathways, highlighting the relevance of the network connectivity [15]. At a first glance, it would be reasonable to assume that a difference in the connectivity of the hVf and dVf neurons existed, i.e., that they were affected by different synapses. We found that the main difference in their connectivity lied in the number of connections from LGN, with dVf neurons receiving almost twice as much as hVf neurons (Tables S2 and S3). In fact, many hVf neurons were not directly connected to LGN, resulting in a zero weighted in-degree (Fig. S4*A*). This finding was consistent with the fact that LGN projects mainly to L4[25], while projections to L2/3 are relatively weak in both weight and connection probability [26].

The analysis of synapses between E2/3 neurons showed that these neurons tended to connect with others of the same class with greater probability (Fig. S4*B*) and strength (Fig. S4*C*). This suggested that dVf neurons exhibited a *rich club* structure, which was also found in models of cortical functional networks [21], and which may facilitate information propagation throughout the column [25]. Similar results have been found in fosGFP expressing layer 2/3 pyramidal cells of the primary somatosensory cortex, which presented an elevated spontaneous activity compared to other pyramidal neurons. Besides, paired-cell recordings showed that these highly responsive neurons had a greater likelihood of being connected to each other [25, 27].

However, due to the relative scarcity of neurons belonging to each of these E2/3 classes and their relatively weak synapses, the total synaptic weight between the different E2/3 classes was negligible compared to, for example, the synaptic weight with E4 neurons (Figs. S4*D-E*). Hence, E2/3 neurons did not substantially influence the particular response of other E2/3 neurons in a direct manner.

In our analysis we also found that the in-degree and out-degree of both classes were quite similar with respect to V1 recurrent connections (Table S2). In particular, dVf and hVf neurons displayed a similar distribution of presynaptic neurons (Fig. S4*D*). Moreover, dVf neurons had slightly stronger incoming recurrent synapses as compared to hVf neurons (Table S3). This occurred primarily due to the contribution of L4 excitatory neurons (Fig. S4*E*), as well as increased LGN input (due to a greater number of LGN connections) (Table S3 and Fig. S4*A*).

Interestingly, a recent experimental study suggested that positive and negative prediction-error neurons in L2/3 map to different transcriptomically defined cell types [28]. If that is correct, then one can expect the above mentioned connectivity differences between these classes – namely, prominent differences in inputs to the different classes of L2/3 neurons from, e.g., L4, but less difference in recurrent connections in L2/3. In this case also the genetic labels of these different classes can help experimentalists to test our modeling predictions regarding connectivity.

The connectivity rules driving the recurrent synapses of the Billeh model (see Methods for details) may introduce some differences regarding the spatial distribution of dVf, hVf and unclassified neurons. To address this, we first studied how individual E2/3 neurons were arranged in layer 2/3. Interestingly, we found that all dVf, hVf and unclassified neurons were equally distributed within the radius and depth of the model (Fig. S5*A*).

To investigate the homogeneity of the spatial distribution of E2/3 neurons and to explore whether neurons of the same class, i.e. dVf, hVf or unclassified, tended to form spatial clusters, we determined for each class the Ripley’s *K* function [29] as a function of the search radius (Fig. S5*B*). The values of *K*(t)/A for the dVf and hVf neurons did not reflect, at least clearly, preferences for grouping neurons with similar functional behaviour. Therefore, although some differences from the random distribution were observed, these were very small, and thus we found no reason to believe that there were substantial preferences in proximity between the dVf and hVf neurons.

### Dynamical differences in E2/3 classes

The effect of the network connectivity was shown to play a role in the emergence of dVf and hVf neurons. However, to gain a more complete understanding of the mechanism that allows the formation of error neurons, it is necessary to analyze the dynamics of the network. To this end, we initially determined the effective synaptic weight, which depends on both the synaptic weight and the activity of the presynaptic neuron, for each presynaptic-postsynaptic class pair (see Methods for details) (Figs. 3 and S6). We found that dVf neurons experienced a large increase in both their recurrent excitatory inputs (mainly from excitatory neurons in L4) and BU inputs (Fig. 3*A*). This excitatory presynaptic activity overcame the increased inhibitory presynaptic activity, mainly from L2/3 and L4 Pvalb populations, resulting in the depolarization of these neurons. In contrast, hVf neurons experienced a slight increase in excitation received from LGN and L4 neurons which was not sufficient to overcome the inhibition originating primarily from L2/3 and L4 Pvalb interneurons, leading to the hyperpolarization of these neurons (Fig. 3*A*). Somewhere between these two classes of neurons were the unclassified neurons, where inhibition originating from Pvalb interneurons was balanced by excitation originating from layer 4 excitatory neurons and LGN input (Fig. 3*B*). Consequently, the different behaviours of the E2/3 neurons were a direct consequence of the excitation received from the BU input and L4 excitatory neurons competing with inhibition from L2/3 and L4 Pvalb interneurons. Experimental evidence in primary somatosensory cortex also revealed that subset of highly active L2/3 excitatory neurons received a significantly larger excitatory input from layer 4 compared to their non-active neighbours, highlighting the relevance of network wiring for the formation of these subsets of neurons [25]. Therefore, a clear understanding of the mechanism leading to the segregation of dVf and hVf neurons can only be obtained by a comprehensive dynamic study of their main sources of presynaptic current.

**Fig. 3.**
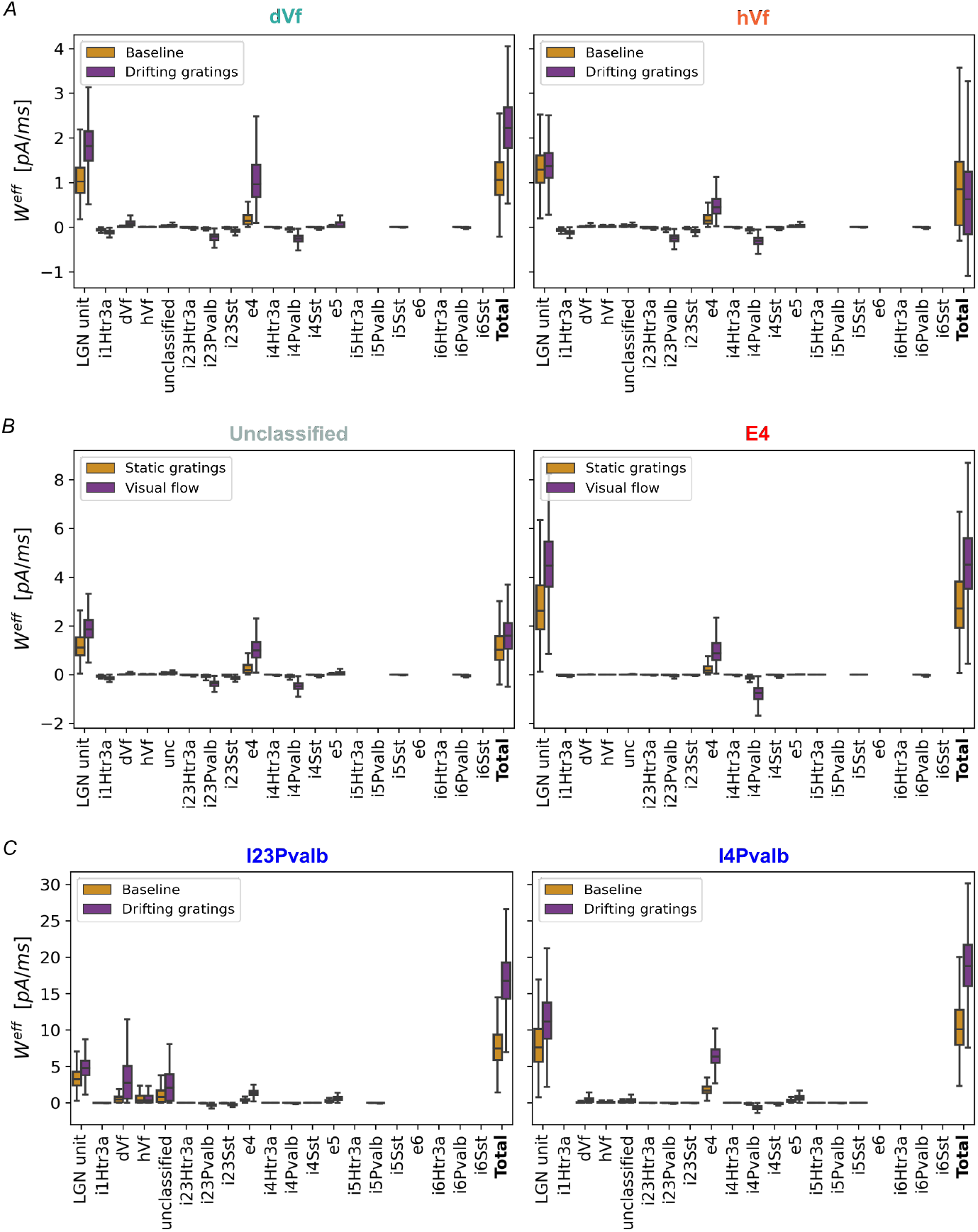
Dynamic origins of E2/3 hyperpolarizing and depolarizing responses to onset of visual flow. *A*, Effective synaptic input weight for dVf and hVf neurons. Bars represent the effective weight 500ms before (orange) and 500ms after (purple) the onset of visual flow. *B*, As in A, but for layer 2/3 unclassified neurons and layer 4 excitatory neurons. *C*, As in A, but for Pvalb neurons in layers 2/3 and 4.

#### L4 excitatory neurons

These neurons acted as amplifiers of the BU excitation (Fig. 3*B*). Notably, this population primarily suffered from increased BU excitation, which triggered a positive feedback loop within the population. This result was consistent with the literature, as the L4 excitatory neuron circuit appears to be located upstream in the local excitatory network and is assumed to be specialized for sensory processing [13].

#### L4 Pvalb interneurons

Similar to their excitatory counterparts, the activity of the L4 Pvalb interneurons increased as a result of the increased activity of both LGN and L4 excitatory neurons (Fig.*3C*). Consequently, they converted the BU-driven excitatory input into an inhibitory input that entered the L2/3 circuitry.

#### L2/3 Pvalb interneurons

This population was mainly responsible for the emergence of dVf and hVf classes of neurons. Their dynamic analysis (Fig. 3*C*) revealed that these neurons were strongly depolarized (Table S1). Although these interneurons also received excitatory input from LGN and neurons in other layers, the main change in their presynaptic activity was caused by dVf neurons. Thus, the activation of dVf neurons drove the depolarization of L2/3 Pvalb interneurons. A detailed analysis of the connectivity between layer 2/3 inhibitory Pvalb neurons and their excitatory counterparts revealed that the former did not connect preferentially with any of the latter classes, i.e., dVf, hVf, or unclassified (Fig. S7*A*) when the total synaptic weight was divided by the number of presynaptic neurons.

Our results also showed that other L2/3 inhibitory-neuron subclasses (Htr3a and Sst), which depolarized as a result of the onset of visual flow (Fig. S6 and Table S1), did not contribute significantly to the dynamics of dVf and hVf neurons (Fig. 3*A*). Experimental evidence suggests that Htr3a and Sst neurons do not receive significant excitation from LGN [26]. However, the presence of a TD input could significantly change the importance of these neuronal populations, as suggested by other computational models explaining the emergence of error neurons [14, 15].

Finally, from the analysis of the model dynamics, it can be seen that the layer E5/6 depolarization is triggered primarily by the input from LGN and L4 excitatory neurons, along with recurrent intralayer excitation (Fig. S6*C*). Thus, the lack of hyperpolarizing responses at L5/6 was the result of insufficient BU-driven recurrent inhibition. Consequently, there was no multi-pathway excitatory-inhibitory balance in recurrent currents for the E5/6 neurons, one of the conditions for the formation of error neurons (Table S1).

### Effect of feature selectivity on the E2/3 classes

GIven the “like-to-like” connectivity rules followed in the construction of the V1 model, it is possible that dVf and hVf neurons have differences also in their direction tuning, since the Billeh model allows for orientation and direction tuning through local, recurrent connections [24]. The preferred direction of stimulus motion of each neuron, which was assigned during model design, varied between the different E2/3 neurons (Fig. S8*A*). For example, dVf neurons tended to prefer vertical gratings that moved horizontally. In contrast, hVf and unclassified neurons exhibited a slight preference for horizontal gratings that moved vertically.

Previously, we classified E2/3 neurons as either dVf or hVf based on their particular response to the sudden horizontal drift of vertical gratings at 2 Hz. Since the dVf and hVf neurons changed their activity when the gratings suddenly started drifting, it is possible that hVf neurons preferred static gratings over the drifting gratings and the contrary for the dVf neurons, as suggested by experimental studies [11]. Thus, we increased the speed of grating drift to 8 Hz and reclassified the E2/3 neurons according to their response to the new stimulus and compared both classifications. We found that neuron assignments to dVf and hVf classes were not significantly affected (Fig. S8*B* and *C*), contrary to what was found in PE neurons, where changes in visual flow speed explained the increased responses of error neurons [4]. This indicates a specific limitation in the Billeh model when processing different temporal dynamics of visual stimuli.

Our analysis revealed that dVf neurons exhibited a particular preference for the gratings orientation (Fig. S8*A*). Could this mean that the particular behavior of a dVf or hVf neuron depended on the orientation of the stimulus? To tackle this question we considered a stimulus consisting of horizontal gratings moving vertically at 2 Hz, to explore the effect of orientation tuning. In this case, the classification of E2/3 neurons changed drastically from the initial classification, suggesting that the two-way behaviour of these neurons was strongly affected by the gratings orientation. This result was the expected since the Billeh model support orientation selectivity. It is worth mentioning that similar results were found for orthogonal gratings in cortical functional network models [21].

We also analyzed the case of having vertical gratings moving at 2 Hz in the opposite direction to the original stimulus (Fig. S8*B* and *C*). It can be seen that the different input currents had not undergone significant changes with respect to the original stimulus. In addition, the reclassification of E2/3 neurons presented a high correlation with the original ones. Consequently, the dynamic behavior of these neurons does not appear to be affected by the stimulus movement direction.

Finally, a stimulus analogous to the original one was considered, but interchanging the periods of visual flow and static gratings. Here, the response of the network was evaluated under a sudden stop of the gratings movement. As expected, the behaviour of dVf and hVf neurons was reversed: hVf neurons depolarized and dVf neurons hyperpolarized (Fig. S8B and C), similar to what was found experimentally [5], but, in our case, in the absence of a TD input.

### Direction selectivity vs perturbation sensitivity

Given the relevance of gratings orientation in the emergence of the dVf and hVf neuron classes, we wondered whether there existed neurons that behaved similarly regardless of grating direction of motion. If the answer was affirmative, it would have been an indication that these neurons responded mainly to the “perturbation” of the visual flow. This type of behaviour was experimentally found in V1 neurons [11]. Until now, E2/3 neurons were classified as dVf and hVf based on their response to horizontally drifting gratings. We then reclassified E2/3 neurons based on their average input current responses to the onset of visual flow in a wide range of movement directions (0° : 315° : 45°, where 0º represented vertical gratings moving to the right and 90º represented horizontal gratings moving downwards), as we originally did with the horizontally drifting vertical gratings (see Methods for details). This analysis revealed that a subset of dVf (241/2948) and hVf (519/3363) neurons maintained their characteristic response regardless the direction of the gratings, i.e., this subset of dVf/hVf neurons depolarized/hyperpolarized for every gratings direction (Fig. S9*A*). Therefore, we identified these subsets of E2/3 neurons as **perturbation-sensitive neurons**, meaning that they responded significantly to the sudden changes in visual flow regardless of its direction of movement. Furthermore, we found that the magnitude of the response was similar for any visual flow direction, with a slight preference for the front-to-back direction (Fig. S9*A*), consistent with experimental observations [31]. If these perturbation-sensitive neurons were actually encoding a sudden change in visual flow during an open-loop scenario, they would be perfect candidates for prediction PE neurons when a locomotion-related signal was introduced into the model. In particular, dVf neurons were identified as candidates for pPE neurons, and hVf neurons for nPE neurons.

Moreover, the consistent responses of this subset of perturbation-sensitive neurons during visual flow across the various grating orientations and drift directions might suggest that hVf neurons would prefer a slower visual flow speed, e.g. static gratings, while dVf neurons would prefer faster visual flow speeds, thus responding positively to an increase in visual flow speed. To explore this possibility in detail, the responses of these neurons to the onset of visual flow were compared at various visual flow temporal frequencies (1 : 9 : 1*Hz*) while maintaining the vertical orientation of the gratings. The preferred frequency for each neuron in this orientation was then determined as the frequency at which the dVf/hVf neuron received the highest/lowest input current during visual flow. Interestingly, we found that dVf and hVf neurons exhibited a fairly similar distribution of preferred frequencies for this particular orientation (Fig. S9*B*), preferring both of them temporal frequencies close to 5Hz. Thus, up to this temporal frequency, the higher the frequency, the greater the characteristic response of these neurons, similar to what the experimental evidence suggests [11].

On the other hand, a subset of dVf (892/2948) and hVf (503/3363) neurons only responded in a particular direction and behaved as unclassified in all other direction, so we identified them as **direction-selective** neurons.

Finally, to further test whether layer 2/3 excitatory perturbation-sensitive neurons encoded a sudden change in visual flow, we presented the model with black-to-white full-field flash and tracked the responses of E2/3 neurons (Fig. 4). Interestingly, dVf/hVf perturbation-sensitive neurons exhibited a sudden increase/decrease in their input current due to the full-field flash, regardless of whether the full-field flash was white or black (Fig. *4C-D*). This response faded after a hundreds of milliseconds with neuron’s membrane potential returning to their baseline values. In comparison, direction-selective neurons presented a significantly different response, behaving both dVf and hVf direction-selective neurons in a similar way (Fig. 4*C-D*). This behaviour confirmed that perturbation-selective neurons acted as general detectors of changes in the visual flow, while direction-selective neurons focused primarily in detecting the movement in a particular direction.

**Fig. 4.**
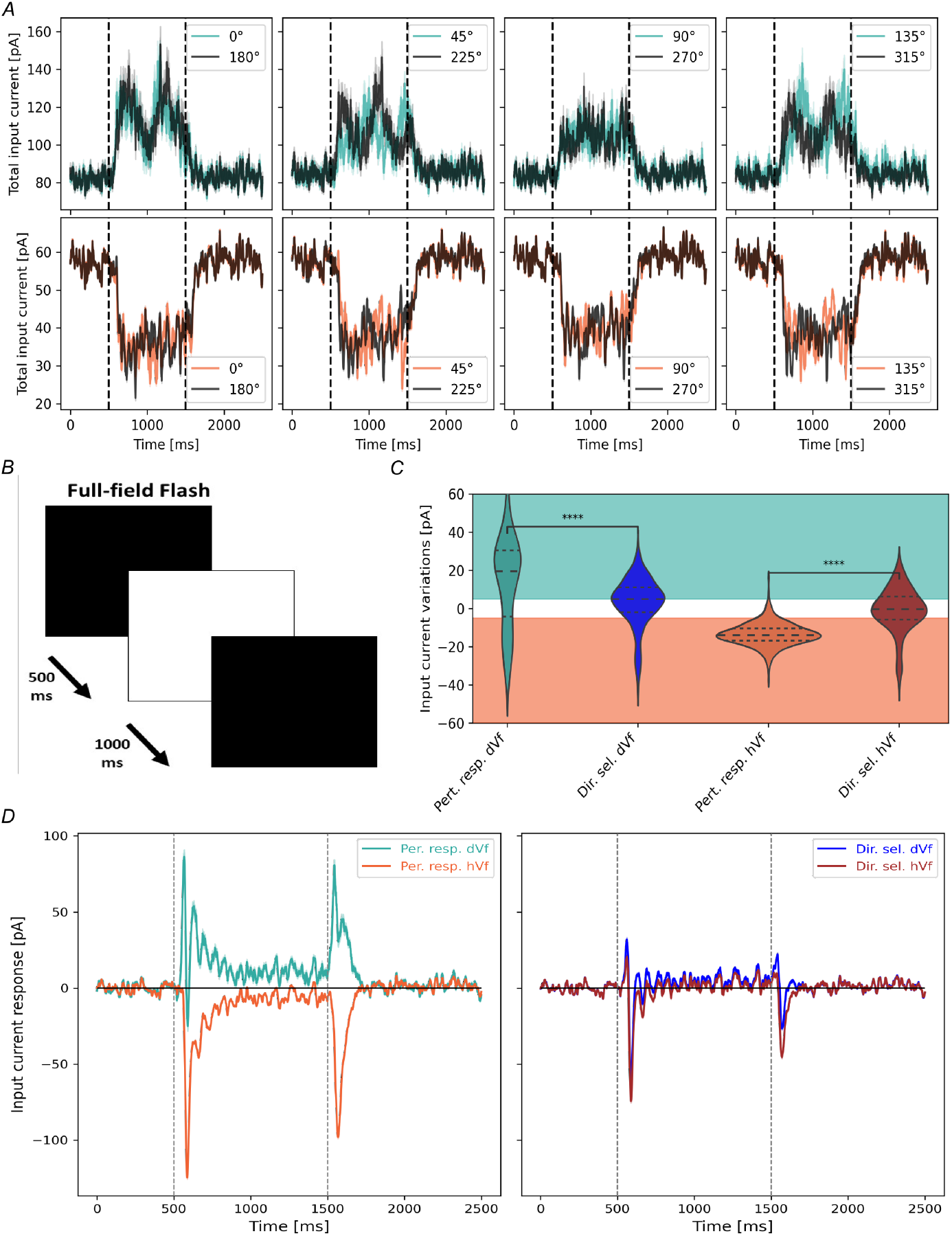
A subset of E2/3 neurons behave as perturbation-sensitive neurons. *A*, Average input current traces for dVf (top) and hVf (bottom) perturbation-sensitive neurons and different grating directions. *B*, Schematics of the full-field flashes. *C*, Probability distributions of the average input current responses to the full field flash for different E2/3 neurons subsets (Dir. sel. dVf vs Pert. resp. dVf, *t* = 6.78, *p* = 7.3 × 10^-11^; Dir. sel. hVf vs Pert. resp. hVf, *t* = −22.1, *p* =1.2 × 10^-83^, Welch’s t test). Turquoise and orange shades indicate regions of dVf and hVf classification. *D*, Trace of the average input current response of E2/3 dVf (turquoise) and hVf (orange) perturbation-sensitive neurons (left) and E2/3 dVf (blue) and hVf (brown) direction selective neurons (right) to the full-field flash. Vertical dotted lines indicate the full-field flash. Shaded areas indicate SEM.

## Discussion

Experimental studies demonstrated the presence of error neurons in L2/3 that report positive and negative errors between predictions of visual flow based on motor signals and the actually seen visual flow [5]. Subsequent work showed that such error neurons can also be detected in the absence of motor signals, suggesting that these error neurons rather report changes in visual flow [11]. These results were obtained even if the perturbation was played back to a stationary animal (in this particular case, smaller responses were elicited). In this study we showed that error neuron-type responses can be obtained using the Billeh model for V1 in the absence of a sensorimotor expectation. More precisely, we found that two functional classes of E2/3 neurons (dVf and hVf) can be found, one of which would report positive errors and the other one negative errors (Fig. 2). This was found despite the fact that, in terms of model composition and wiring, the Billeh model only assumes one model of E2/3 neurons. The dVf and hVf functional classes of neurons that we found behave similarly to depolarizing (dMM) and hyperpolarizing (hMM) mismatch neurons found in [5], where visual flow was uncoupled from locomotion speed, and perturbations consisted of horizontal gratings drifting at random times.

Notably, these two classes of error neurons emerged because of differences in both their synaptic connectivity and their effective connectivity, especially with inhibitory neurons. While dVf neurons received stronger connections from the LGN and other excitatory neurons (mainly from L4), hVf neurons received much fewer connections from the LGN, showing a lack of BU-driven excitatory input (Fig. S4*A*). Therefore, the BU input did not have a direct functional influence on most hVf neurons. Furthermore, dVf neurons also received stronger inputs from layer 4 excitatory neurons than hVf neurons did (Fig. 3*A*). These differences in structural connectivity represent a prediction that could be experimentally tested if these two classes of error neurons were identified.

The dynamic analysis of the network also revealed that the visually driven excitation from LGN and E4 overcame the recurrent inhibition of the network in dVf neurons, resulting in their BU-driven depolarization. Interestingly, dVf neurons played another important role: they depolarized L2/3 Pvalb interneurons. In contrast, hVf neurons received weaker inputs from E4 neurons, rendering them unable to overcome the recurrent inhibition from L2/3 Pvalb interneurons. Consequently, these neurons underwent BU-driven hyperpolarization. Unclassified neurons exhibited a balance between recurrent inhibition and BU-driven excitation targeting these neurons, resulting in little or no response to visual flow perturbation (Table S1).

Recent publications on cortical microcircuit modeling have revealed interesting conditions for the emergence of prediction error neurons [14, 15]. They considered a particular circuit of BU and TD connections and emphasized that while the distribution of BU and TD inputs to excitatory cells was well studied, the distribution between different types of cortical inhibitory interneurons was less understood and probably diverse. Therefore, various input configurations led to different circuits for the error neurons. In our case, the thalamocortical connectivity of the Billeh model, which backs up on experimental studies [26, 32, 33], suggests that BU almost exclusively targets excitatory and Pvalb neurons in all layers and Htr3a neurons in L1. This led us to the circuitry shown in Fig. 5, where dVf neurons indirectly inhibited hVf neurons during the onset of visual flow, by activating L2/3 Pvalb interneurons. This circuit is reminiscent of a *competition by common inhibition* between dVf and hVf neurons, where the former use the pool of Parvalbumin neurons to suppress the activity of the latter [34]. Further evidence for this hypothesis can be seen in Fig. 2*B*, where the firing rate of dVf neurons is anti-correlated with the firing rate of hVf neurons. This outlines a modeling prediction of a connectivity motif, which our work suggests to be important for maintaining the dVf and hVf functional classes, and which can potentially be observed experimentally if one can correlate functional properties of L2/3 neurons with their local connectivity.

**Fig. 5.**
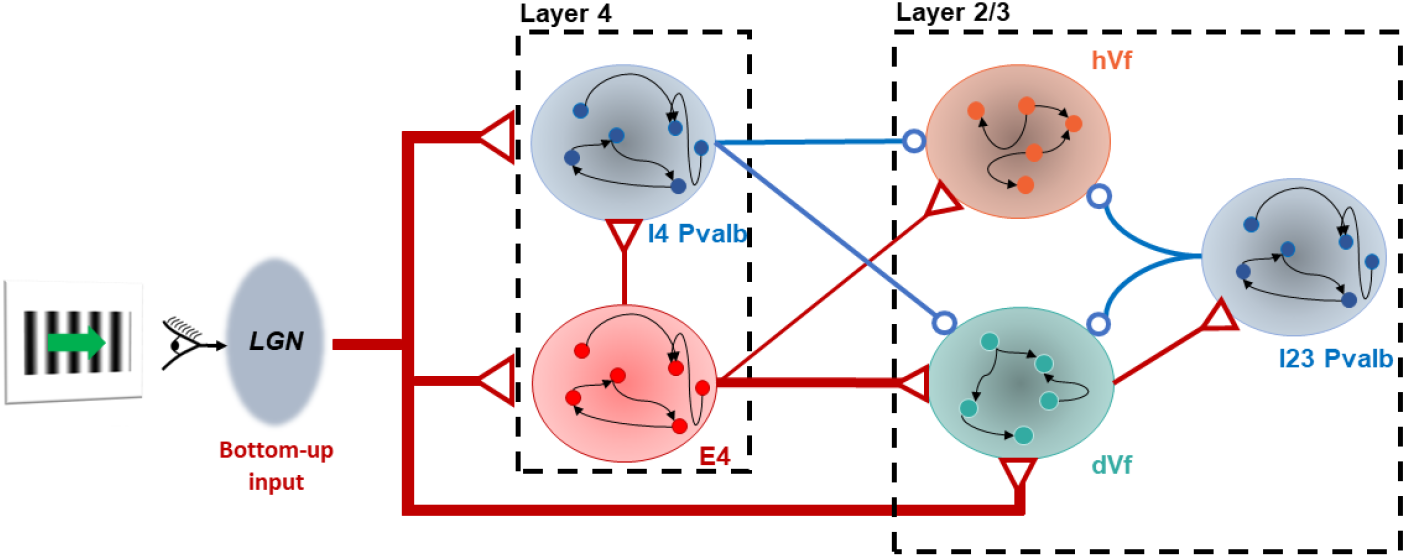
Dynamic network analysis reveals crucial L2/3 inhibitory pathways. Diagram illustrating the effect that the BU excitatory input of LGN has on the segregation of hVf and dVf neurons. Arrows represent variations in effective synaptic weights due to the onset of visual flow. Red arrows indicate excitatory projections, while blue arrows indicate inhibitory projections. Based on dynamic analysis, only neuronal populations that play an important role in the E2/3 neuronal behavioural split are shown.

Previous studies had not considered the pronounced feature selectivity of V1 cells. We found that dVf neurons identified for a certain orientation of a drifting grating tended to prefer that orientation relative to others, whereas unclassified neurons tended to prefer perpendicular gratings (Fig. S8*A*). Moreover, when the vertical orientation of the gratings was changed to a horizontal orientation (Fig. S8), some dVf and hVf neurons behaved as unclassified neurons, while others continued to behave in the same way. Furthermore, a subset of dVf and hVf neurons maintained their characteristic response for all tested drift directions, suggesting that they only respond to visual flow perturbation. This was confirmed through full-field flash stimulations (Fig. 4). Consequently, this subset of dVf and hVf neurons could be associated with the positive and negative perturbation-sensitive neurons found in [11].

Finally, it is worth mentioning that, by construction, the version of the Billeh model that we used is biologically plausible and relevant, despite it describes neuronal cells as point neurons. However, recent modeling studies suggest that prediction errors are computed not in separate neurons, but locally in dendritic compartments [14, 35]. Hence, studying the roles of dendritic computations in producing the effects we described above would be a potentially fruitful area for future work.

In conclusion, the emergence of two classes of neurons that depolarize and hyperpolarize in response to a sudden change in visual flow can be explained by orientation selectivity of neurons and their connectivity, thus being an inherent property of the network architecture. This suggests that error neurons can arise naturally from diverse and realistic neural connections with feature selectivity, as we found in the Billeh model, where no particular training to identify visual errors has been performed. In particular, even in the absence of a TD input, the local V1 circuitry can generate the error neuron-type responses in layer 2/3. It is then expected that this mechanism might work together with additional mechanisms furnished by the TD inputs to enhance or modify these error responses.

## Methods

### Data-driven cortical laminar microcircuit model

Modeling cortical microcircuits represents a difficult task due to the large number of genetically, morphologically, and electrophysiologically different neuron types present in the mammalian neocortex. Furthermore, there is limited insight into the particular connectivity between different pairs of neuron types. However, intensive research in recent years [36–38] led to a detailed cortical microcircuit model for area V1 of mice [24] (Figs. 1*A* and 6). Since this model has been shown to successfully reproduce many computational properties reported in real V1 circuits [24], we implemented its point-neuron version to gain insight into the dynamic organization of the network.

**Fig. 6.**
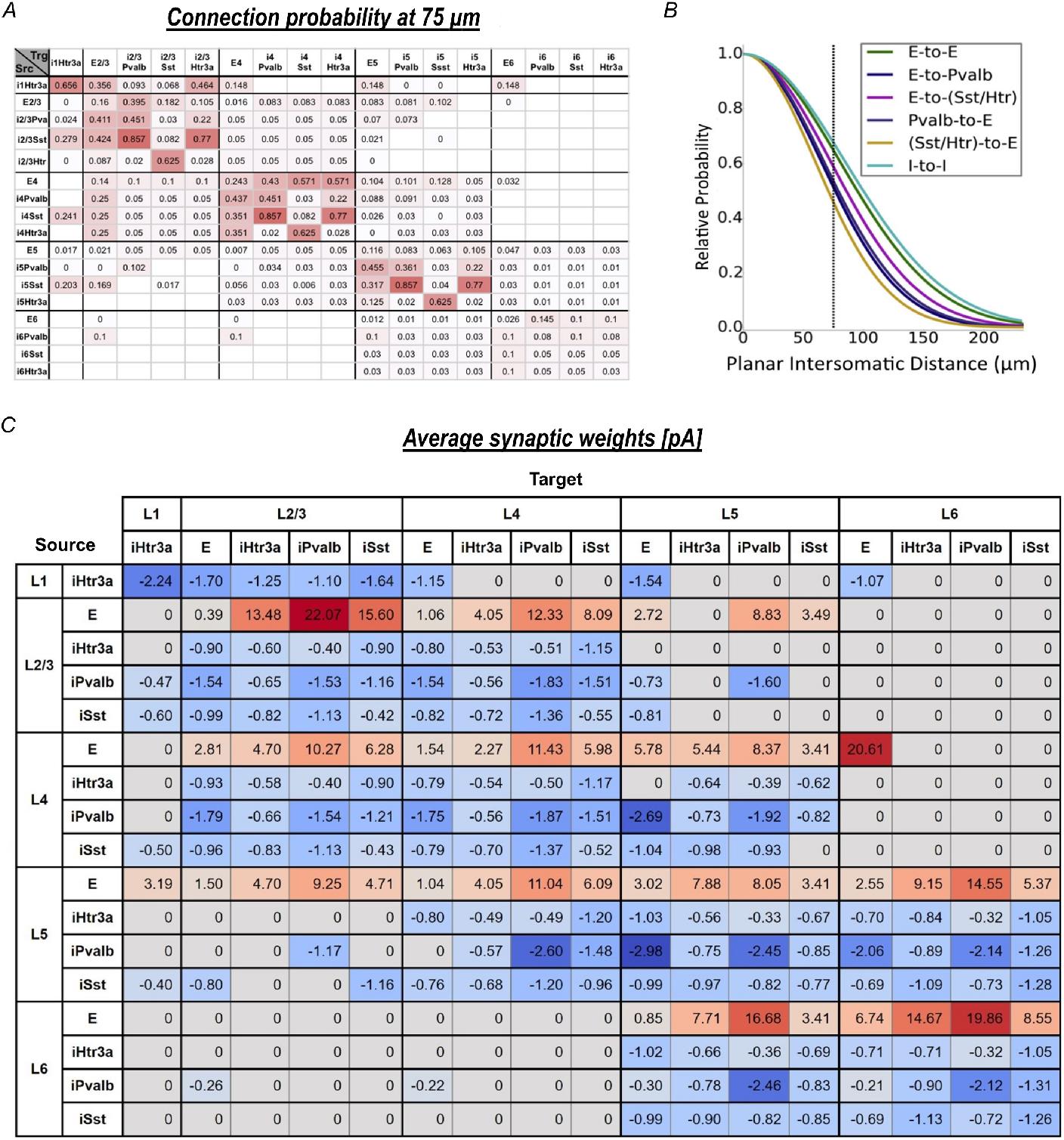
Overview of the data-driven cortical laminar microcircuit model of [24]. *A*, Base connection probabilities at 75*μm* intersomatic distance depending on presynaptic (Src) and postsynaptic (Trg) neuron types. White grid cells indicate unknown values, which are assumed to be zero in the model. *B*, Scaling of the base connection probabilities shown in *B* as a function of the somatic distance for the different types of connections. The probability of a synaptic connection is obtained by multiplying the base connection probability for the presynaptic and postsynaptic neurons with this scaling function. *C*, Average synaptic weights depending on presynaptic (Source) and postsynaptic (Target) neuron types. Gray grid cells indicate unknown values, which are assumed to be zero in the model.

The cortical microcircuit point-neuron version of the model represented a 845μm-radius patch of area V1. The patch was composed by a “core region”, an internal region of 400*μm*-radius, surrounded by an annulus that avoided possible border artifacts. The model was complemented with a LGN module that generated action potentials for arbitrary visual stimuli (Fig. 1*A*). Point neurons were described using a generalized LIF model, named *GLIF*_3_, which contained two internal variables describing the after-spike currents (see below). We focused in this study on the dynamics of the “core region” of the model.

The model contained 230,924 point neurons, 51,978 of which were in the core. Neurons were divided into one class of excitatory and three subclasses of inhibitory: Htr3a-positive (Htr3a) (embracing VIP neurons), Parvalbumin (Pvalb) and somatostatin (Sst). This was the case for layers L2/3, L4, L5, and L6, while L1 contained only Htr3a inhibitory neurons (the precise abundance of each neuron class is shown in Fig. 1*A*). This resulted in 17 different data-based neuron types subdivided into 111 separate models based on individual neuron response profiles from the Allen Cell Types Database [39]. Both the abundance of each type of neuron in each layer and the probability of connection between neurons were based on experimental data. In particular, the base connection probabilities for any pair of neurons of the 17 subclasses, at a horizontal intersomatic distance of 75*μm*, were provided in [24] and are reproduced in the table of Fig. 6*A*. White cells in the grid indicate unknown values in their connections and were taken as zero in the model. The connection probabilities between pairs of cells were scaled by a function that decayed exponentially with the horizontal distance between the two GLIF neurons (Fig. 6*B*).

An important feature of the model is that it assumes that all E2/3 neurons are identical. Consequently, the input to these neurons was solely responsible for the different dynamic behaviors that they exhibited.

#### Leaky Integrate-and-Fire model with after-spike currents (GLIF3)

This neuron model is a modified version of the stereotypical leaky integrate-and-fire (LIF) neuron in which the long-term effects of ion channels opening are modeled as additional *after-spike currents* (ASC) *I_j_*, which has predefined time scales (*k_j_*), a multiplicative constant (*R_j_*), which is typically set to 1, and a constant (*A_j_*) that is added to each current after a spike [40]. Therefore, the evolution equation of the membrane potential is given in [24] as

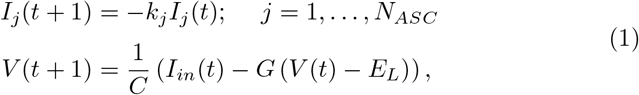

where *N_ASC_* = 2 is the number of after-spike currents, *C* is the neuron capacitance, *G* is the membrane conductance, *E_L_* is the resting membrane potential, and *I_in_* is the input current which can be decomposed as

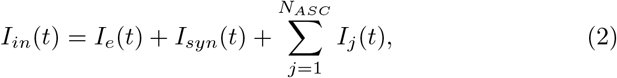

where *I_e_* is the time-dependent external current and *I_syn_* is the post-synaptic current. The update rule, which is applied if *V* (*t*) > *θ*_∞_, is:

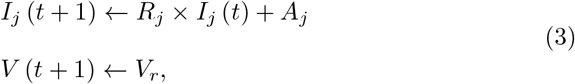

where *V_r_* is the reset membrane potential. After a spike, the voltage of a neuron remains at its reset value for a certain refractory period (*t_ref_*), which depending on the neuron model takes values between 2 ms and 8 ms. The time step in all the simulations was set to 1 ms. This neuron model was fitted by maximizing the likelihood that a model neuron with intrinsic noise exactly reproduces the spike train observed in real neurons [40].

#### Synaptic features

In the Billeh model, the dynamic of the synapses is described by the following function:

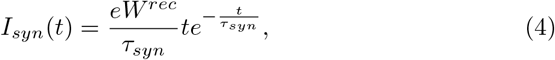

where *I_syn_* is the postsynaptic current, *τ_syn_* is the synaptic time constant, and *W^rec^* is the synaptic weight. The constants *τ_syn_* took values 5.5 ms for excitatory-to-excitatory (EE) synapses, 8.5 ms for inhibitory-to-excitatory (IE) synapses, 2.8 ms for excitatory-to-inhibitory synapses, and 5.8 ms for inhibitory-to-inhibitory (II) synapses. Axonal delays were distributed over the range [1, 4] ms, which was extracted from Fig. 4*E* of [24] and converted to integers since the integration step is 1 ms. Additionally, postsynaptic weights were established taking into account the planar intersomatic distance between neurons, functional rules assuming similarity in preferred direction angle, and distance in retinotopic visual space, resulting in the synaptic weights shown in Fig. 6*C* (see [24] for details). Three main feature-related functional rules were taken into account:

1. The Billeh model accounts for “like-to-like” preferences within excitatory-to-excitatory synapses in L2/3 excitatory circuitry, meaning that neurons that prefer similar direction of motion are preferentially connected, these connections also being stronger [24, 41]. In addition, similar connectivity rules applied to the strength (but not the probability) of other synaptic classes.
2. For E-to-E classes only, a decrease of synaptic strength with distance in retinotopic visual space between source and target neurons, projected to the target neuron’s preferred direction.
3. A retinotopic correction in E-to-E synaptic weights to avoid assymetric retinotopic magnification.

In total there were 70,139,111 synapses and the probability of connection between two randomly chosen neurons in the network was 0.263%.

#### External inputs

The LGN in the thalamus is known to project retinal inputs to V1 via excitatory projections, which we refer to as bottom-up input [42]. In the Billeh model, the LGN was modeled by a sample of 17,400 LGN units, each one represented by a spatio-temporally separable filter. These units operated with images and movies in the visual space as inputs and returned time series of instantaneous firing rate, which were converted to spike trains using a Poisson process, as output. Each LGN unit was sampled from 14 subclasses that approximated the diversity observed in the mice LGN. Furthermore, LGN units selectively innervated only excitatory and Pvalb neurons in L2/3-L6 and Htr3a neurons in L1, with L4 being the main target of these projections, as discussed by [13]. Therefore, the current contribution of the LGN to each GLIF neuron was given by

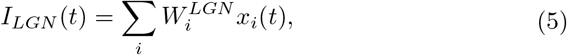

where 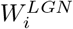 is the postsynaptic weight of the LGN unit *i*, the variable *x_i_*(*t*) takes the value 1 if at time *t* the LGN unit fires, and 0 if it does not, and the sum extends over all LGN units. A second external excitatory input source was included. It consisted of background noise (BKG) from neurons that were not considered in the circuit. This noise was described by a single Poisson source firing at 1kHz and was injected into all V1 neurons. This noise source had different synaptic weights depending on the target neuron class and contributed to the external current as *I_BKG_*. Thus, the external current received by each GLIF neuron was given by

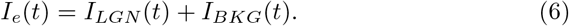

### Visual stimulus

#### Drifting gratings

The visual stimuli used in the simulations were similar to those used in the experiments [5, 11] and consisted of a full-field sinusoidal grating drifting in a given direction with spatial frequency 0.04cycles/° and 0.8 contrast. In our case, for each simulation we included an initial 500 ms duration presentation of the static grating. After that period, the previously static grating abruptly started drifting (perturbation) for a time of 1000 ms. After this period, a static grating was presented again for 1000 ms.

#### Full field flashes

In the latest part of our work we used a full-field black image for a time of 500 ms. Then a period consisting of a full-field white image was presented for a duration of 1000 ms. After this period we performed a new black image presentation for 1000 ms.

#### Analysis of the results

For every simulation, we recorded the spikes, membrane potential, and input currents for each neuron at each integration step. However, due to the strong reset of the membrane potential after a spike, the membrane potential did not represent a useful variable to study the network dynamics. Instead, we found that the currents injected into each neuron were the appropriate parameter to characterize the dynamics of the neurons. Baseline values were determined through their mean value during the 500*ms* prior to the visual flow onset. Responses to the visual flow onset were obtained by subtracting the baseline value. Additionally, firing rates were sampled at 60*Hz* and smoothed with a 150 ms Gaussian filter.

#### Determination of the rheobase

For each cell model we determined its *rheobase, θ_rheo_*, as the minimum amount of current injected into the soma, for a rectangular shaped current injection, that elicited a spike. For that purpose we injected rectangular pulses of 1 s duration into the cell model. We started at 1 pA and subsequently increased it by 0.01 pA interleaved by 1 s resting periods until a spike took place. Once the cell started to fire, the corresponding injected current was identified as its rheobase (Fig. S1 illustrates the mentioned process).

#### Classification of E2/3 neurons

Neurons were identified as depolarized with visual flow (dVf) if the average value of the input current variation, ΔI_in_, given by

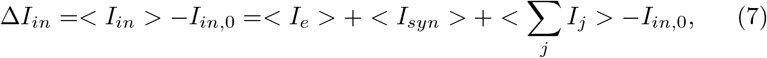

exceeded a certain threshold, *θ_d_*. In the equation *I*_*in*,0_ is the baseline value, *I_e_* is the external current (6), *I_syn_* is the postsynaptic current arriving from other V1 neurons (4), and *I_j_* represent the *j*-th ASC (3). Average over 20 trials and 1000 ms duration of visual flow were taken. Similarly, neurons were classified as hyperpolarized with visual flow (hVf) if the average value of the input current response was lower than a certain value *θ_h_*. Otherwise, neurons were labeled as unclassified. Since each cell type had different characteristics we took as upper threshold 0.05 times the cell rheobase, and as lower threshold −0.05 times the cell rheobase. Although these threshold values were arbitrary, we found that our results were robust over a range of threshold values.

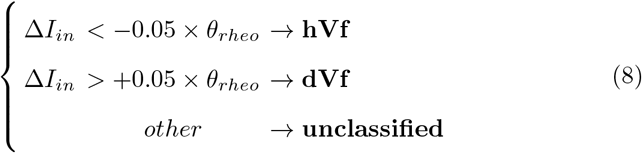

#### Ripley’s *K* function

To investigate the homogeneity of the spatial distribution of E2/3 neurons and to explore whether neurons of the same class, i.e. dVf, hVf or unclassified, tended to form spatial clusters, we determined for each class the Ripley’s *K* function [29] defined as

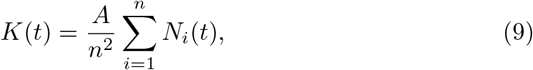

where *A* is the area of layer 2/3, *t* is the search radius, the index *i* runs over the cells of a given class, and *n* is the total number of cells in the class. *N_i_*(*t*) stands for the number of cells of the type *i* within a intersomatic planar distance *t*. When the search area falls partially outside of the V1 column, we apply a weighting factor based on the ratio of the search area that falls within V1 to the total search area. The effects of edge corrections are more important for large *t* because large search circles are more likely to be outside the V1 column, leading to an underestimation of Ripley’s K real value. This indicator measures the spatial heterogeneity of a certain type of cells. The baseline for random homogeneous systems (or null model) is a growth of type *k*(*t*) = *πt^2^* until the system edge is reached. If the value of *K*(*t*) is above the random curve for a certain search radius *t*, it implies that the network is spatially clustered at that scale compared to a null model.

#### Effective synaptic weight

We defined the *effective synaptic weight* between a presynaptic neuron *i* and a postsynaptic neuron *j* as

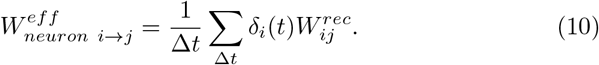

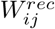 represents the synaptic weight (see Eq. (4)), *δ_i_*(*t*) took the value 1 if at time *t* the presynaptic neuron fired and 0 otherwise, and the sum extended over the period Δ*t* (ms). To determine the *effective synaptic weight* from neuron class *i* (presynaptic) to neuron *j* (postsynaptic), we considered the contribution of each presynaptic neuron of class *i* to neuron *j*,

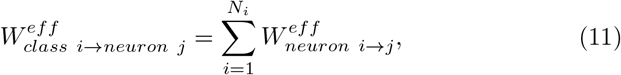

where *N_i_* represent the abundance of neurons of class *i*.

### Statistics

For each simulation 20 trials were computed (the first one was discarded for later analysis) and the time traces of each measured quantity were averaged. For comparisons of mean values involving more than two classes (e.g., hVf, dVf and unclassified L2/3 neurons), a one-way ANOVA test was first performed. In case of a significant result (*p* < 0.01), two-sided Welch’s t-tests were used to determine if there was a significant difference between the mean values of the populations; NS=not significant, \**p* < 0.05, * * *p* < 0.01, * * \**p* < 0.001, * * * * *p* < 0.0001. If only two distributions were compared (e.g., E2/3 and E5/6), a two-sided Welch test was performed directly.

Box-and-whisker plots are drawn so that the box represents the interquartile range and the median, and the whiskers represent the 10*^th^* and 90*^th^* percentiles.

## Data availability

All data is generated by the simulation code (see Code availability statement below).

## Code availability

All simulation code used for this paper is available on GitHub (https://github.com/JavierGalvan9/Billeh_model_error_neurons). The model was implement on Tensorflow [43] software and simulated in GPU.

## Acknowledgments

This work is partially supported by the Severo Ochoa and Maria de Maeztu Program for Centers and Units of Excellence in R&D, grant MDM-2017-0711 funded by MCIN/AEI/10.13039/501100011033.

## Declarations

### Authors contributions

J.G.F.: writing-original draft, software, formal analysis, writing—review & editing. F.S.: Software. J.J.R.: Conceptualization, writing—review & editing. A.A.: Conceptualization, writing—review & editing, supervision. W.M.: Conceptualization, investigation, writing—review & editing, supervision. C.R.M.: Conceptualization, investigation, writing—review & editing, coordination of the research.

## Additional information

An animated representation of the model can be seen in https://s10.gifyu.com/images/model_representation.gif.

More details on the biological features of E2/3 neurons can be found at the Allen Brain Atlas http://celltypes.brain-map.org/experiment/electrophysiology/487661754.

An example of the visual stimuli can be checked at https://www.youtube.com/watch?v=wKstDX3xw0k.

## Supplementary material

### Determination of the cells model rheobase

**Fig. S1.**
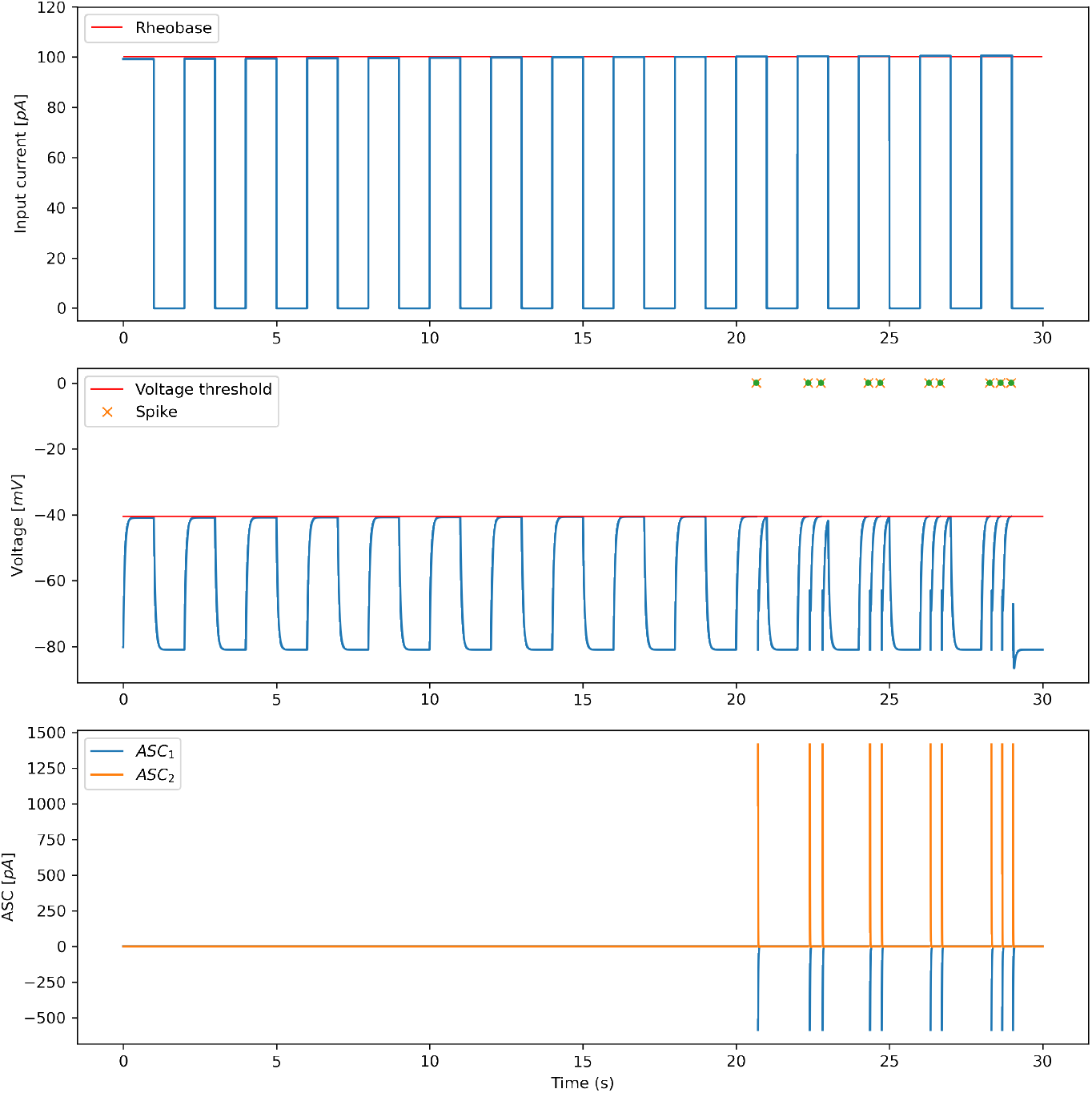
Illustration of the procedure followed for the determination of cell models rheobase. Top: Injected current into the cell soma consisting of increasingly larger current steps interleaved by resting periods. Red line represents the identified rheobase. Middle: Membrane voltage of the cell model as a response to the injected current. Red line represents the voltage threshold in the model. Bottom: After-spike currents of the model. Sharp vertical lines indicate the presence of a spike.

### Sample responses from E2/3 neurons

**Fig. S2.**
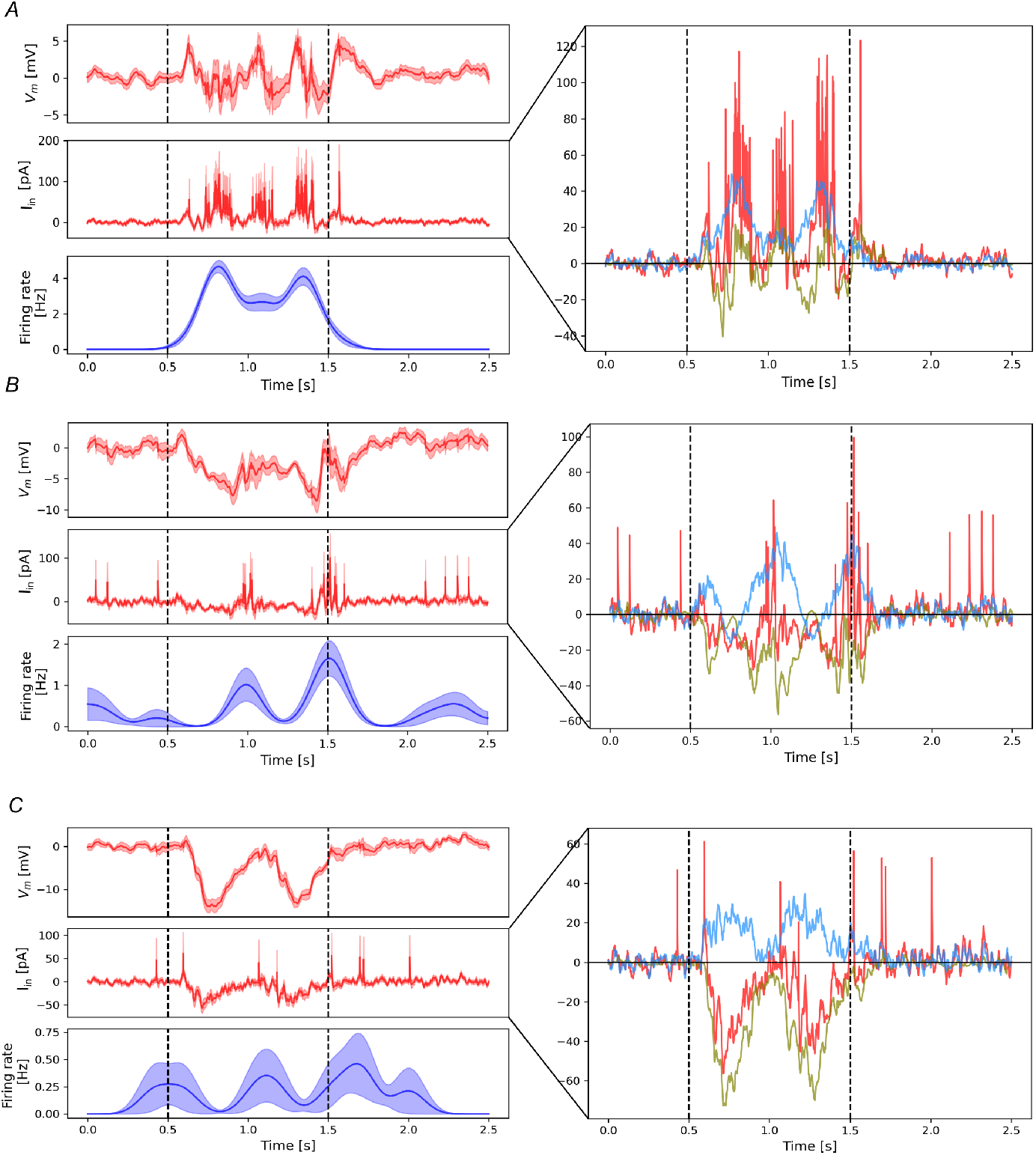
Responses to the sudden onset of visual flow for one neuron of each E2/3 class. *A*, Left: Trace of the neuron’s membrane voltage response (top), input current response (mid) and firing rate (bottom) for a dVf sample neuron. The mean input current during visual flow (blue horizontal line) and the classification thresholds (black dots) are shown. Right: Trace of the responses of the different input current components for a sample dVf neuron: total input current (red), recurrent current (olive) and BU current (blue). Vertical dashed lines indicate the visual flow period. *B*, As in *A*, but for an unclassified neuron. *C*, As in *A*, but for a hVf neuron.

**Table S1.** Input currents for different neuron populations.

| Population | Class | Abundance | Baseline currents [$pA$] | | | Visual flow currents [$pA$] | | |
| --- | --- | --- | --- | --- | --- | --- | --- | --- |
|  |  |  | Recurrent | BU | Total | Recurrent | BU | Total |
| E2/3 | dVf | 23.2% | 0.32±0.06 | 14.8±0.1 | **73.2±0.1** | 2.5±0.2 | 25.3±0.2 | **84.4±0.2** |
|  | hVf | 26.5% | -0.34±0.06 | 11.6±0.2 | **69.3±0.2** | -9.8±0.1 | 12.2±0.2 | **60.9±0.2** |
|  | unc | 50.3% | -0.10±0.04 | 14.4±0.1 | **72.4±0.1** | -5.1±0.1 | 19.3±0.1 | **72.2±0.1** |
| I2/3 Pvalb | dVf | 100% | 35.9±0.5 | 20.4±0.5 | **72.3±0.8** | 91.0±0.7 | 29.1±0.7 | **133±1** |
| I2/3 Htr3a | dVf | 70% | 6.8±0.1 | 0 | **25.1±0.2** | 19.2±0.2 | 0 | **35.1±0.3** |
|  | unc | 30% | 6.3±0.1 | 0 | **28.4±0.4** | 13.9±0.2 | 0 | **35.9±0.5** |
| I2/3 Sst | dVf | 100% | 15.0±0.2 | 0 | **50.2±0.5** | 39.6±0.4 | 0 | **77.2±0.6** |
| E4 | dVf | 52.4% | 0.2±0.0 | 36.9±0.2 | **35.7±0.2** | -1.1±0.1 | 60.5±0.2 | **51.5±0.2** |
|  | hVf | 10.9% | 1.6±0.1 | 52.1±0.5 | **47.8±0.4** | -5.8±0.2 | 50.4±0.5 | **36.9±0.4** |
|  | unc | 36.7% | 0.3±0.0 | 43.3±0.2 | **41.5±0.2** | -4.1±0.1 | 51.5±0.3 | **42.9±0.2** |
| I4 Pvalb | dVf | 96.1% | 17.9±0.2 | 58.6±0.8 | **72.9±0.7** | 51.3±0.4 | 83.6±0.9 | **118.5±0.7** |
|  | unc | 3.9% | 23±1 | 101±4 | **110±4** | 49±2 | 89±4 | **117±3** |
| E5/6 | dVf | 62.7% | 9.0±0.1 | 13.4±0.1 | **95.7±0.2** | 34.4±0.2 | 19.8±0.1 | **115.6±0.2** |
|  | hVf | 0.3% | 15±3 | 21.2±0.9 | **80±3** | 8±3 | 21.8±0.7 | **71±3** |
|  | unc | 37.0% | 7.2±0.1 | 14.1±0.1 | **93.2±0.2** | 13.6±0.1 | 18.0±0.1 | **99.6±0.2** |

Mean values for the different input current components during the static gratings (baseline) and drifting gratings (visual flow) periods. In some populations a classification of the belonging neurons was made (see Methods for details). The SEM on the sample population and realizations is taken as the error. The ASC and BKG inputs are not shown but do contribute to the total current.

### Baseline input current distributions of E2/3 neurons

**Fig. S3.**
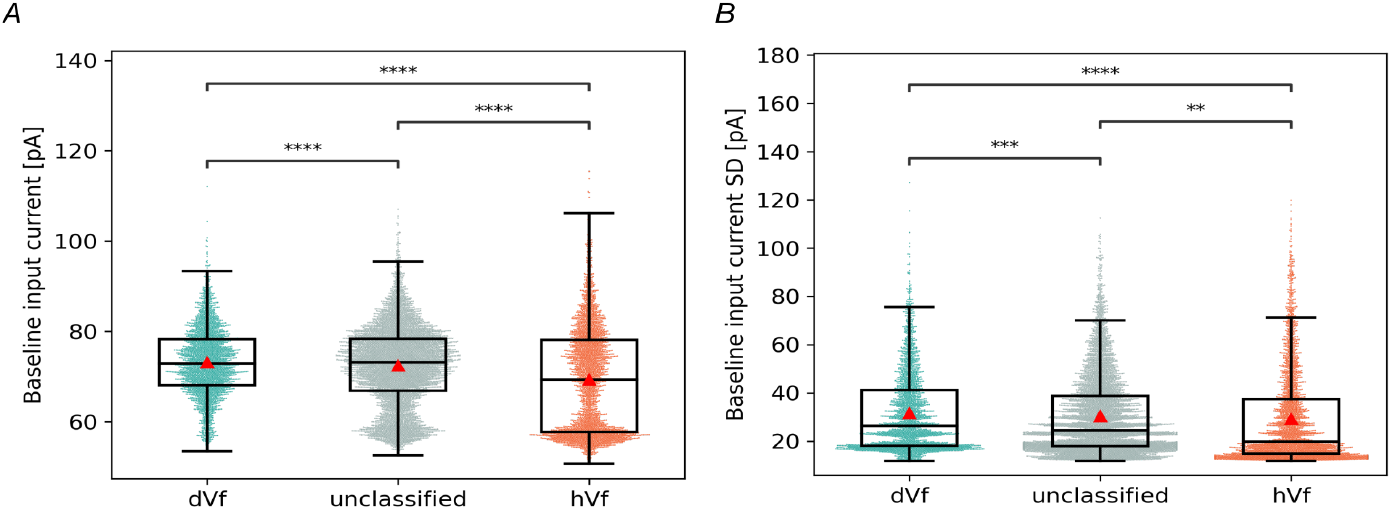
Baseline distributions of E2/3 neurons. *A*, Distribution of baseline input current mean for E2/3 classes (ANOVA: *F* = 162.8, *p* = 1.5 × 10^-70^, *n* = 12689; dVf vs hVf: *t* = 15.72, *p* =1.5 × 10^-54^, *n* = 6311; dVf vs unc: *t* = 4.18, *p* = 2.9 × 10^-5^, *n* = 9326; hVf vs unc: *t* = 13.6, *p* = 2.8 × 10^-41^, *n* = 9741, Welch’s t test). *B*, Distribution of baseline input current standard deviation (SD) for E2/3 classes (ANOVA: *F* = 14.96, *p* = 3.2 × 10^-7^, *n* = 12689; dVf vs hVf: *t* = 5.24, *p* =1.7 × 10^-7^, *n* = 6311; dVf vs unc: *t* = 3.58, *p* = 3.5 × 10^-4^, *n* = 9326; hVf vs unc: *t* = 2.72, *p* = 0.0065, *n* = 9741, Welch’s t test). Red triangles indicate the class mean.

### Connectivity of E2/3 neurons

To better understand the connectivity of the E2/3 classes, it is interesting to analyze the degree and weighted degree of their neurons. In particular, the **in-degree** (*k_in_*) of a neuron is the number of synapses it has where the given neuron is the postsynaptic one. Similarly, the **out-degree** (*k_out_*) of a neuron is the number of synapses it has where the given neuron is the presynaptic one (Table S2). However, since each synaptic connection presents a weight, it is more convenient to consider **weighted degrees**, also called **total synaptic weight**, which follow the same definition as degrees but are weighted according to the weight of each connection (Table S3). Regarding the in-degree, it can be divided into input from external sources (LGN and BKG) and V1 recurrent connections.

**Table S2.** In/Out degrees for E2/3 classes.

| Class | Out-degree ($k_{out}$) | In-degree ($k_{in}$) | | | **Total** |
| --- | --- | --- | --- | --- | --- |
|  |  | Recurrent | LGN | BKG |  |
| dVf | 337.7±0.3 | 469.3±0.4 | 16.5±0.1 | 1.0±0.0 | **486.8±0.4** |
| hVf | 338.1±0.3 | 471.9±0.4 | 8.7±0.1 | 1.0±0.0 | **481.6±0.4** |
| unc | 337.5±0.2 | 469.3±0.3 | 13.2±0.1 | 1.0±0.0 | **483.5±0.3** |

Connections with other V1 neurons (recurrent), thalamus (LGN) and noisy background sources (BKG) are considered. The SEM for each variable is taken as the error.

**Table S3.** Weighted in/out degrees for E2/3 classes.

| Class | Weighted out-degree ($k_{out}$) | Weighted in-degree ($k_{in}$) | | | **Total** |
| --- | --- | --- | --- | --- | --- |
|  |  | Recurrent | LGN | BKG |  |
| dVf | 1067±3 | 404±3 | 297±2 | 4.0±0.0 | **704±3** |
| hVf | 1081±3 | 302±2 | 157±3 | 4.0±0.0 | **463±3** |
| unc | 1074±2 | 353±2 | 237±2 | 4.0±0.0 | **594±2** |

Connections with other V1 neurons (recurrent), thalamus (LGN) and noisy background sources (BKG) are considered. The SEM for each variable is taken as the error.

**Fig. S4.**
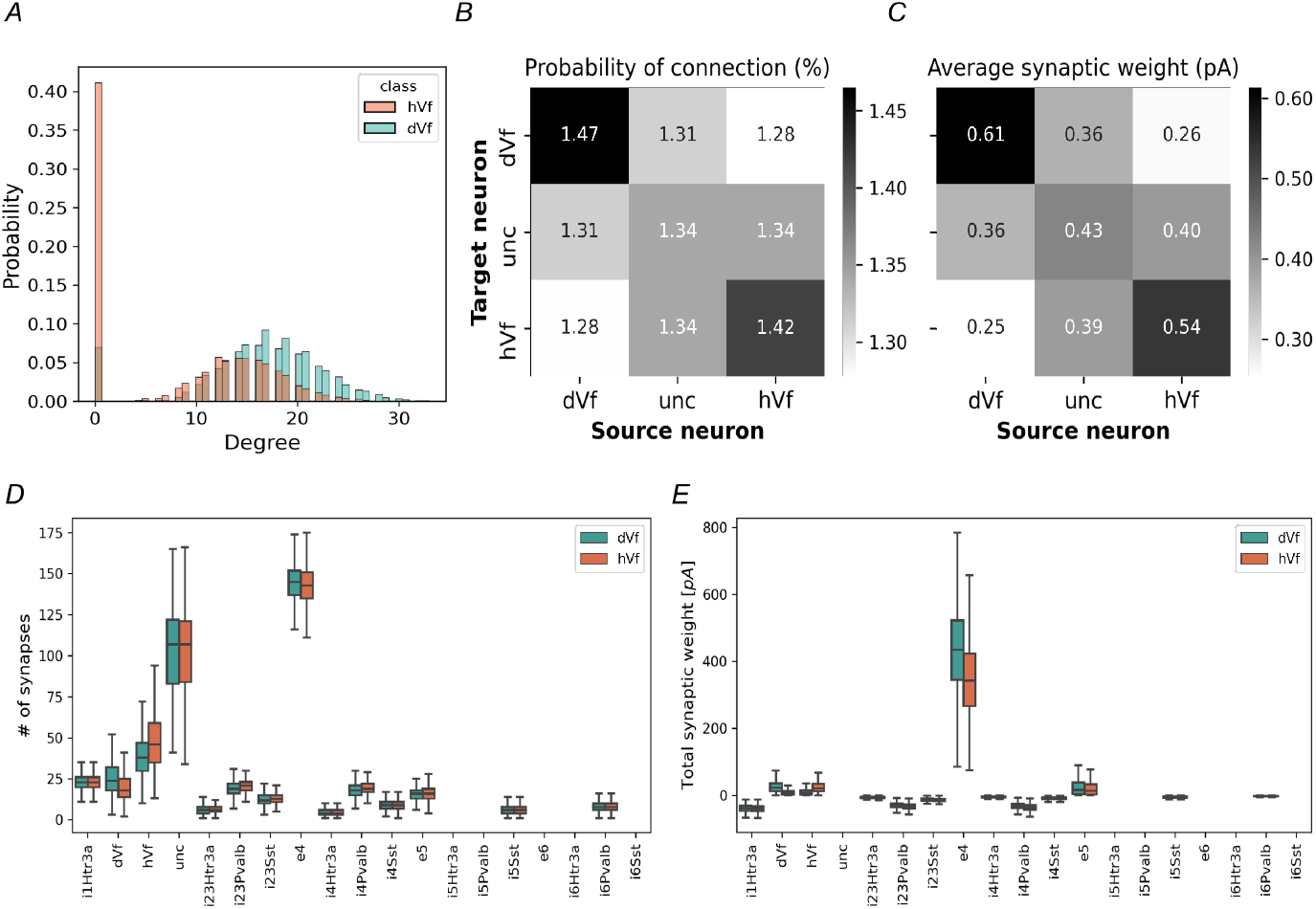
Connectivity of E2/3 neurons. *A*, Distribution of weighted in-degrees from LGN units for hVf (orange) and dVf (torquoise) neuron classes. *B*, Connection probability (%) of two randomly selected E2/3 neurons. Note the symmetry of the matrices, which shows that there is no bias in the source-target order. *C*, Average synaptic weight of a given connection between two randomly selected E2/3 neurons. *D*, Number of synapses with different populations of presynaptic neurons for the hVf (orange) and dVf (torquoise) classes. *E*, Average synaptic weight with different populations of presynaptic neurons for the hVf (orange) and dVf (torquoise) classes.

### Spatial distribution of E2/3 classes

**Fig. S5.**
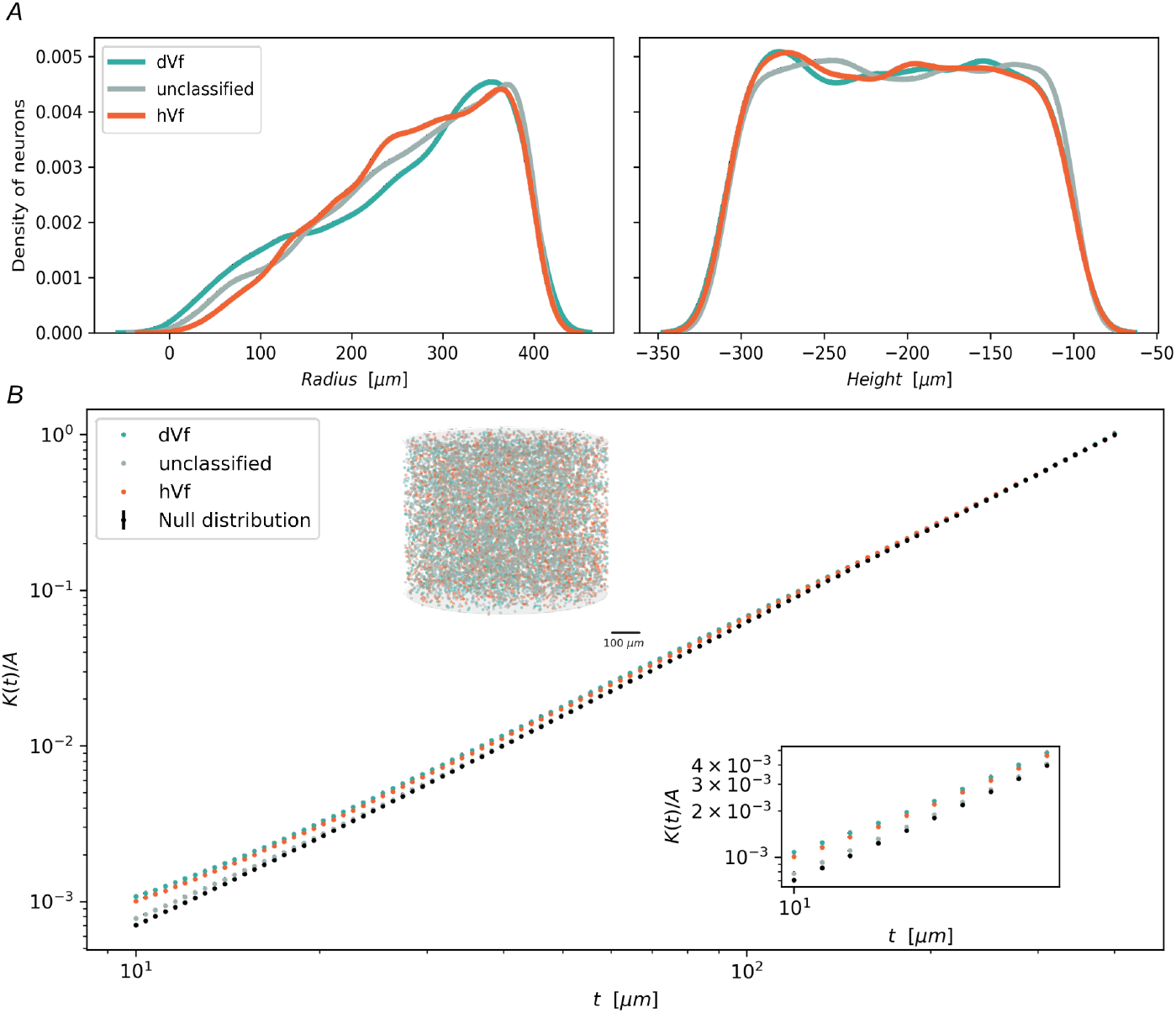
Spatial distribution of dVf, hVf and unclassified neurons. *A*, Density distribution of the distance to the center of the column, which we refer to as radius, (left), and the depth within the layer (right) for dVf, hVf and unclassified neurons. *B*, Ripley’s *K* divided by the layer 2/3 area as a function of the search radius *t*. The colours of the curves and dots represent the results for dVf (turquoise), hVf (orange) and unclassified (gray) neurons. The black points represents the outcome of a random null model averaged over 100 realizations. Also, the E2/3 classes are represented in the V1 cylinder.

### Dynamic weights for different pairs of pre- and postsynaptic neurons

**Fig. S6.**
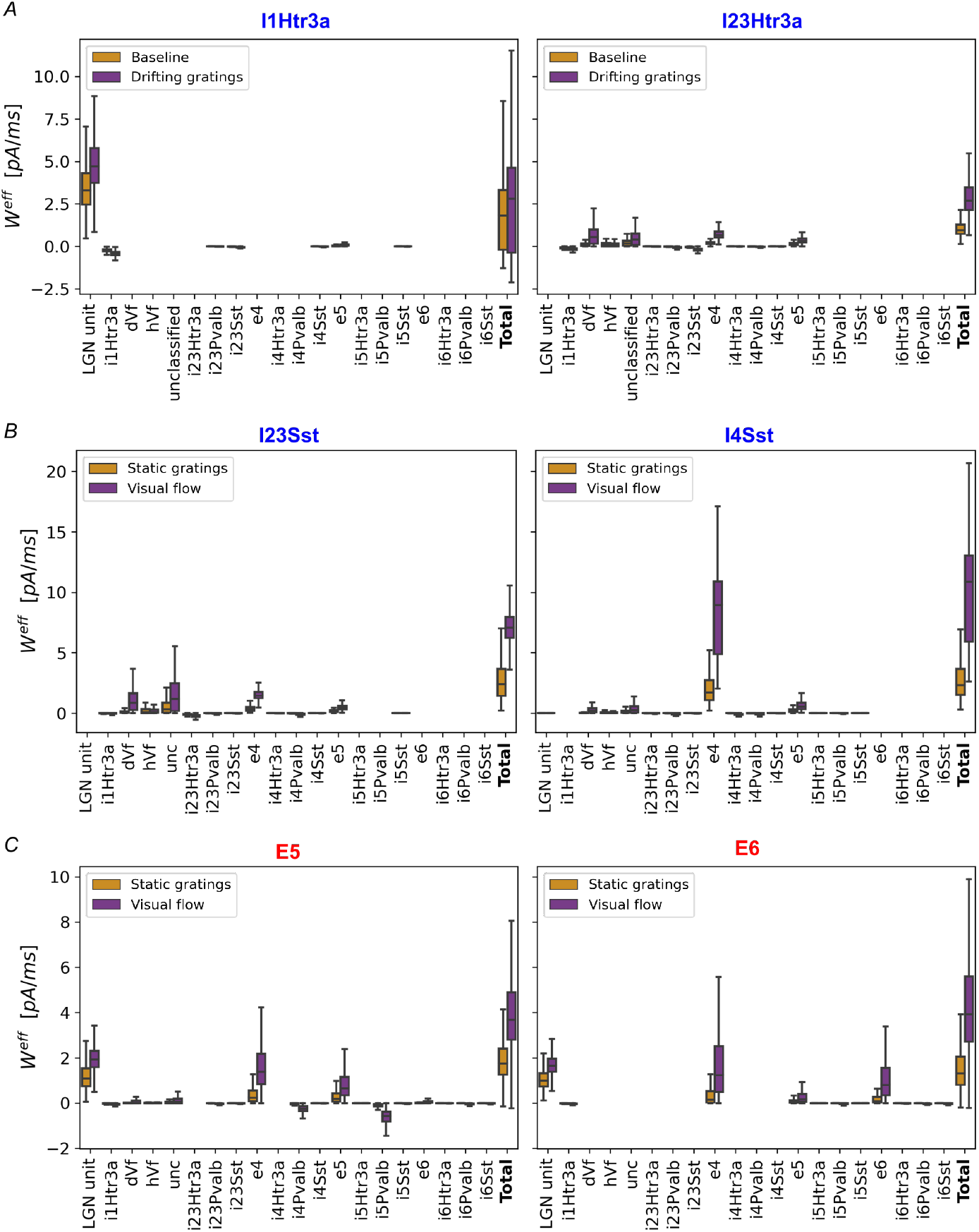
Distributions of effective synaptic weights for different neuronal populations. Effective synaptic weight for various classes of neurons. The bars represent the effective weight 500ms before (orange) and 500ms after (purple) the onset of visual flow.

### Synaptic weights for different pairs of pre- and postsynaptic neurons

**Fig. S7.**
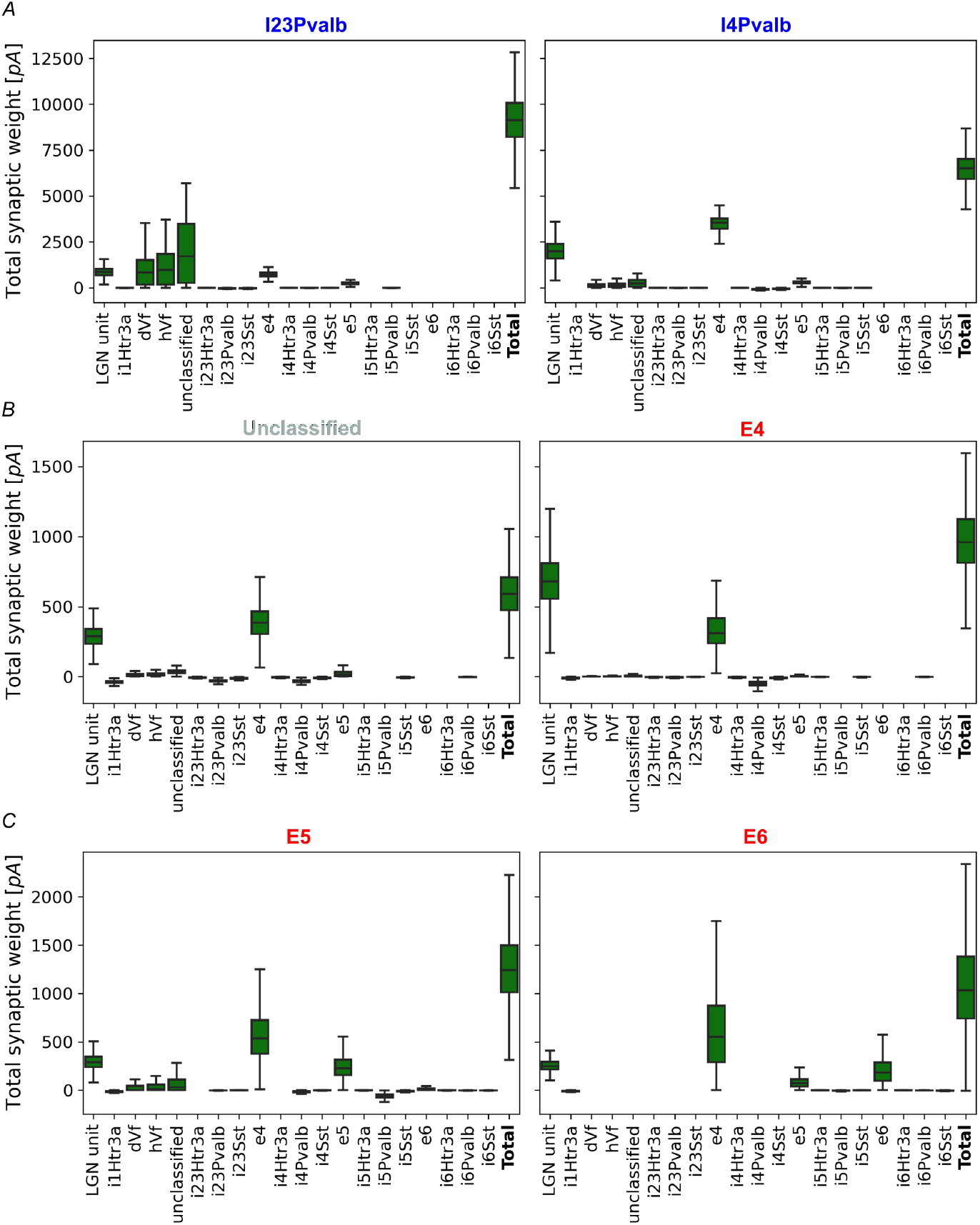
Presynaptic weight distributions for different neuronal populations. Total synaptic weight of several presynaptic populations for: *A*, L2/3 and L4 inhibitory Parvalbumin neurons; *B*, L2/3 unclassified and E4 neurons; and *C*, E5 and E6 neurons.

### Responses of E2/3 neurons to different BU input features

**Fig. S8.**
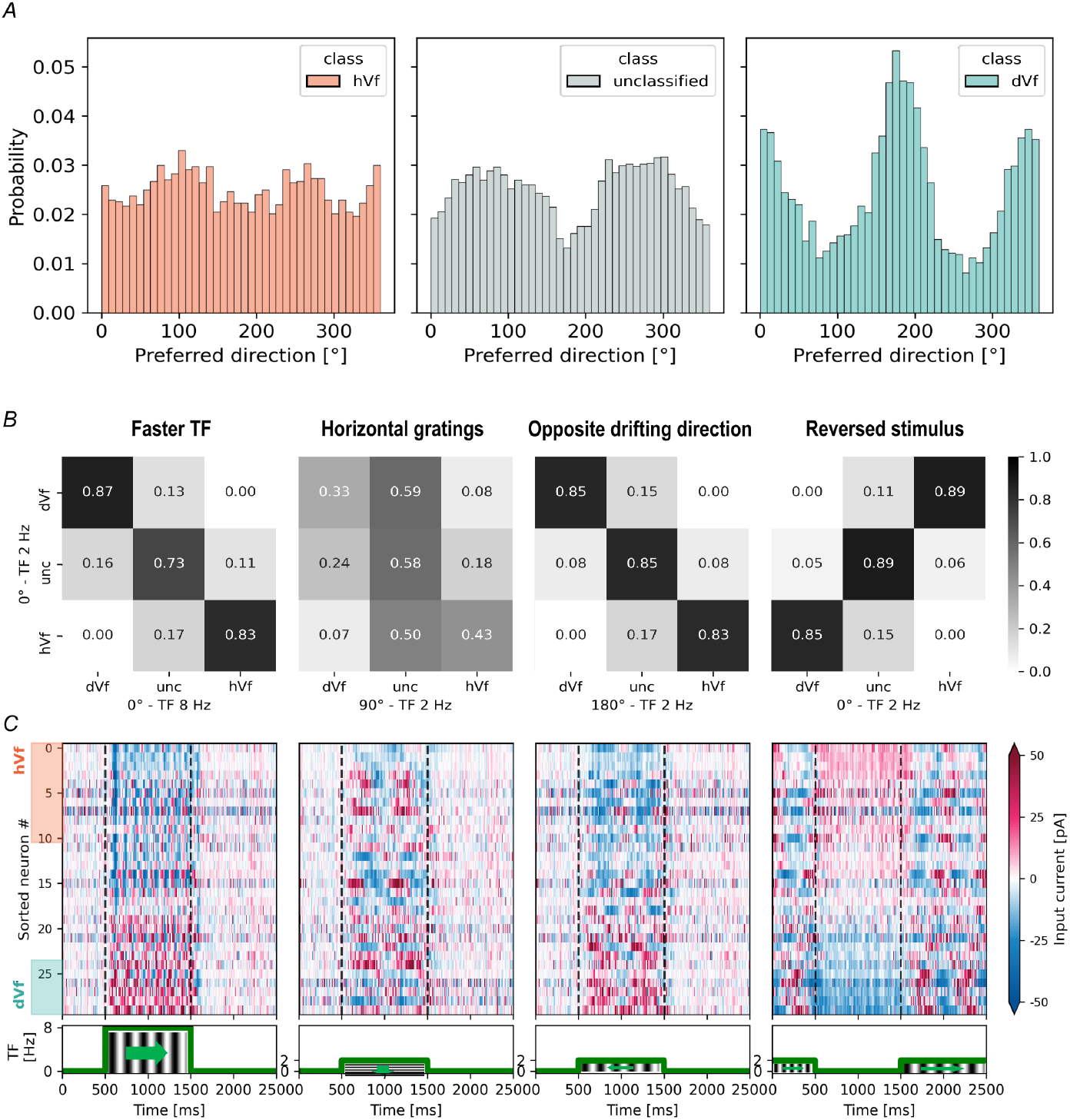
Responses of E2/3 neuron classes to various BU input features. *A*, Distribution of preferred direction of stimulus motion for E2/3 dVf (left), unclassified (middle), and hVf (right) neurons. *B* and *C*, The different visual stimuli consisted of a faster visual flow composed of vertical gratings moving at 8Hz (left), horizontal gratings moving vertically (middle left), a reversed direction of motion (middle right), and a sudden stop of visual flow (right). *B*, Confusion matrices showing the fraction (0-1) of E2/3 neurons classified in each class according to their response to the new stimulus (horizontal axis) compared to the classification of the original experiment (vertical axis). *C*, Heatmap of the average input current responses to the given stimulus for the same neurons shown in Figure 2*A*. TF represents the temporal frequency of the visual flow.

### Characteristics of E2/3 perturbation-responsive neurons

**Fig. S9.**
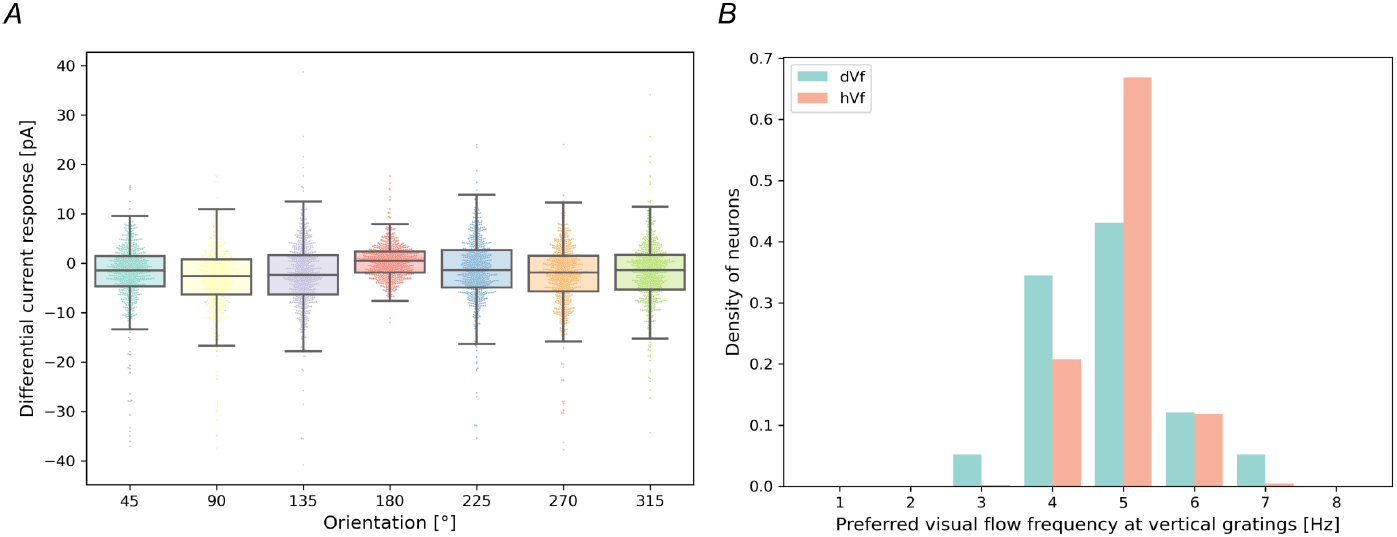
Orientation and frequency preferences of E2/3 perturbation-responsive neurons. *A*, Input current responses in various drift directions normalized to the input current responses to horizontal drift (0 direction). *B*, Distribution of preferred visual flow frequency of dVf (turquoise) and hVf (orange) neurons, and horizontal drift vertical gratings.

### V1 neurons response to full-field flash

**Fig. S10.**
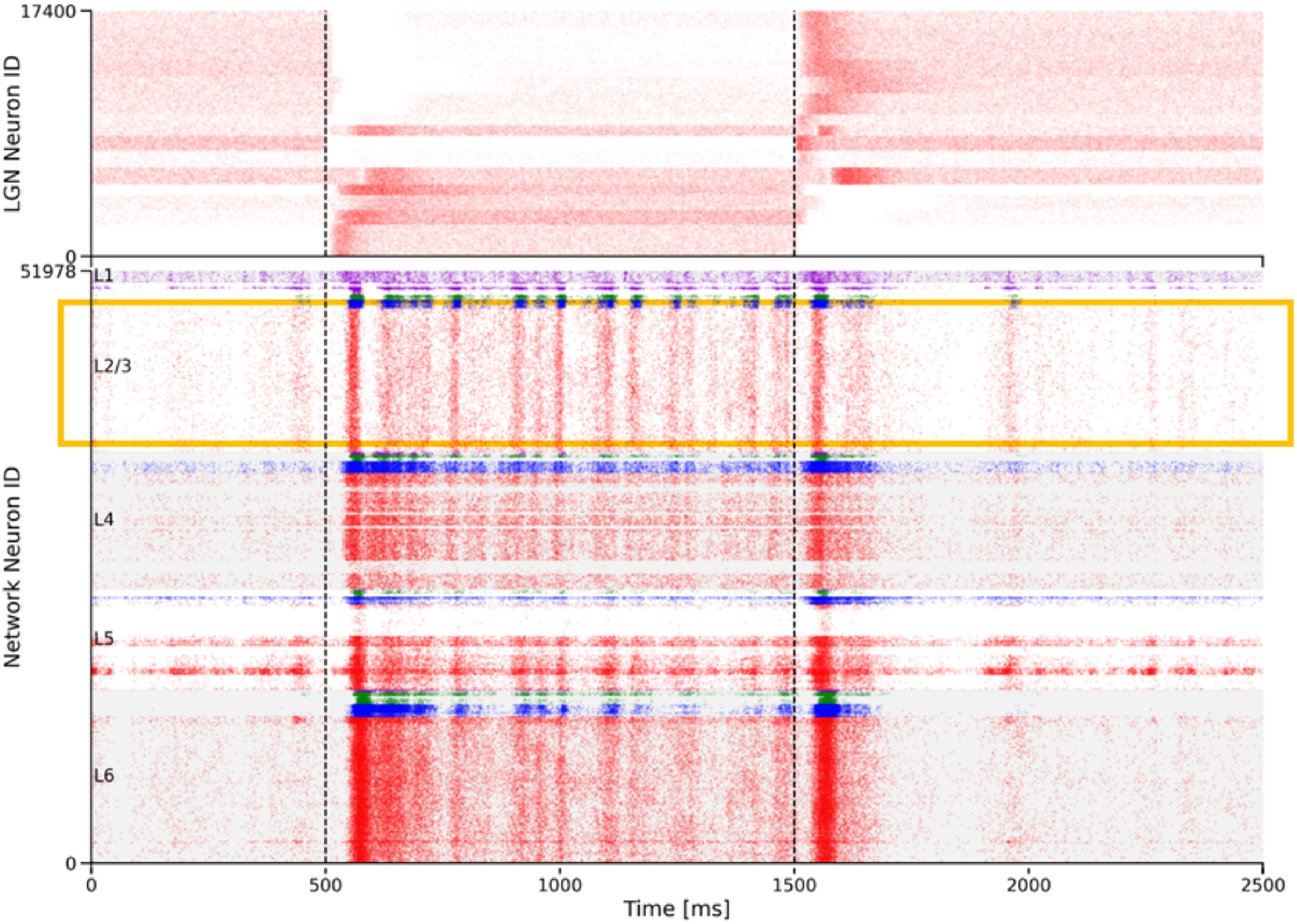
Raster plot of V1 model response to full-field flash stimulus. Top: Raster plot of the spike response of LGN units to full-field flash. Bottom: Laminar raster plot of the spike response of V1 neurons to full-field flash. The colors of the spikes represent the different populations of neurons, following the same palette as in Fig 1. Vertical dashed lines indicate the period of full-field flash.

